# Resolving Emergent Transient Oscillations in Gene Circuits with a Growth-Coupled Model

**DOI:** 10.1101/2025.07.09.663992

**Authors:** Hari R. Namboothiri, Ayush Pandey, Chelsea Y. Hu

## Abstract

Synthetic gene circuits often behave unpredictably in batch cultures, where shifting physiological states are rarely accounted for in conventional models. Here, we find that degradation-tagged protein reporters could exhibit transient oscillatory expression, which standard single-scale models do not capture. We resolve this discrepancy by developing Gene Expression Across Growth Stages (GEAGS), a dual-scale modeling framework that explicitly couples intracellular gene expression to logistic population growth. Using a chemical reaction network (CRN) model with growth-phase-dependent rate-modifying functions, GEAGS accurately reproduces the observed transient oscillations and identifies amino acid recycling and growth-phase transition as key drivers. We reduce the model to an effective form for practical use and demonstrate its adaptability by applying it to layered feedback circuits, resolving long-standing mismatches between model predictions and measured dynamics. These results establish GEAGS as a generalizable platform for predicting emergent behaviors in synthetic gene circuits and underscore the importance of multiscale modeling for robust circuit design in dynamic environments.

**Teaser:** Multiscale modeling reveals how growth and proteolysis-linked recycling cause transient oscillations in synthetic gene circuits.

## Introduction

Synthetic biology has enabled the construction of gene circuits that sense, compute, and actuate within living cells. As the field advances from coarse-grained designs to increasingly sophisticated, multilayered systems, the ability to predict and control circuit behavior over time becomes essential. Dynamical models, which describe how molecular species change over time, are fundamental tools for understanding and engineering these systems. They allow researchers to simulate circuit trajectories, tune design parameters, and anticipate failure modes. This modeling capacity is especially critical when synthetic gene circuits operate in complex or changing environments, where both the molecular dynamics and host physiology evolve. As synthetic biology moves toward real-world applications, from live therapeutics to industrial bioprocesses, accurate and predictive modeling becomes a prerequisite for rational design.

Dynamical models have played a pivotal role in the design and optimization of synthetic gene circuits, particularly in the model organisms such as *Escherichia coli (E. coli)*. Reduced models with lumped parameters representing transcription and translation rates have enabled rational circuit design, improving performance*(1-3*), supporting the implementation of feedback control(*4-6*), and guiding strategies to mitigate resource competition(*7-9*). In parallel, large-scale chemical reaction network (CRN) models have gained traction through the growth of omics datasets and computational tools(*10*). These frameworks have been applied to metabolic pathway engineering(*11, 12*), microbial community modeling(*13*), and whole-cell simulations for high-throughput phenotype prediction(*14*). However, the predictive accuracy of many existing models remains limited by their narrow focus on a single physical scale. Most dynamical models assume constant growth rate when describing intracellular processes, while models at the population scale often omit gene expression dynamics. In practice, growth and gene expression are tightly coupled and vary continuously over time, creating complex multiscale interactions that current models rarely capture. These interactions can give rise to emergent dynamics, where system-level behaviors arise that are not predictable by considering molecular or population scales individually. This disconnect complicates efforts to engineer circuits with reliable dynamics, often prolonging the design–build–test–learn (DBTL) cycle and limiting translational success(*15*).

There has been long-standing interest in understanding how gene expression is coupled to cell growth in bacterial systems(*16-18*). Growth dynamics reflect the availability of intracellular resources and the influence of extracellular conditions on global physiological states. For example, ribosome abundance(*19, 20*), RNA polymerase concentration(*20, 21*), tRNA pool size(*20, 22*), and translation elongation rates (*23, 24*) have all been shown to vary systematically with growth rate. These empirical observations have motivated the development of coarse-grained models linking growth and gene expression, grounded in bacterial growth laws and proteome allocation theories(*16, 21, 25, 26*). While these models establish a conceptual and quantitative connection between growth rate and expression capacity, they typically assume constant exponential growth. This assumption limits their utility for modeling gene circuits that operate in dynamical environments. Some studies have introduced growth perturbations into gene expression models(*27*), simulated transitions between growth phases(*28*), or examined host–circuit interactions in specific contexts(*29, 30*). Previous studies have reported functional links between population growth and circuit dynamics, but the results are narrow in scope and lack strong mechanistic support(*4, 31*). A generalizable, modular, and transferable framework spanning molecular and population scales is therefore required to capture multiscale dynamics(*32*).

In this work, we address the gap between gene expression and population growth modeling by developing a dual-scale framework called Gene Expression Across Growth Stages (GEAGS). Our approach was motivated by a consistent discrepancy between single-scale model predictions and experimental data from a simple gene expression cassette encoding a degradation-tagged fluorescent reporter. While conventional models predicted a monotonic, faster response due to increased protein turnover, batch culture experiments instead revealed unexpected transient oscillations, which suggests that key mechanistic details were missing from the model. We hypothesized that growth-dependent resource dynamics play a central role. To investigate this, we constructed a CRN model linking molecular and population scales through three extendable rate-modifying functions (RMFs). This dual-scale model accurately reproduced the observed transient oscillations. Sensitivity and phase portrait analysis identified growth dynamics and amino acid recycling as key contributors to this behavior. To make the framework more accessible for design applications, we derived a reduced six-state model that preserves the essential dynamical features of the full CRN. This effective model distills the full multiscale CRN into a compact system with a small number of interpretable parameters, preserving key growth-dependent features while enabling practical design exploration and tuning. To demonstrate generalizability, we further applied the GEAGS framework to a layered feedback control circuit(*4*). By integrating RMFs into existing effective models, we were able to accurately capture the growth phase-dependent dynamics of all four controller architectures. Together, these results establish GEAGS as an adaptable and extendable modeling tool for synthetic gene circuit design in batch cultures and other dynamical growth contexts.

## Results

### Model-experiment discrepancies reveal hidden multiscale dynamics

Systems design in synthetic biology often relies on minimal models that simplify gene expression into coarse-grained dynamics. For instance, the process of gene transcription and translation of a single gene expression cassette (GEC) can be reduced to two ordinary differential equations (ODEs) with two states: mRNA (*M*) and Protein (*P*), where *P* is the measurable output, usually a fluorescent protein(*32*):

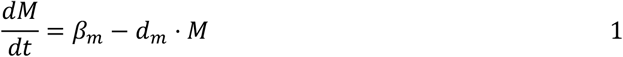

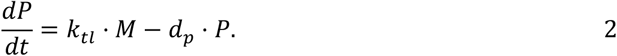

In this system, *β*_*m*_ and *k*_*tl*_ denote the rates of transcription and translation, respectively, while *d*_*m*_ represents the degradation rate of mRNA. The parameter *d*_*p*_ accounts for the combined effects of protein degradation (*d*_deg_) and dilution (*d*_dil_), such that *d*_*p*_ = *d*_deg_ + *d*_dil_. Under conditions of constant cell division, *d*_*p*_ is dominated by dilution (*d*_*dil*_). During exponential growth, *E. coli* cells typically double in 20 to 40 min(*33*), while green fluorescent protein (GFP) variants without degrons have half-lives ≥ 24 hours (*34*), therefore dilution occurs at a considerably faster rate than protein degradation (*d*_*deg*_). A key metric for characterizing gene expression dynamics is the response time, which describes how quickly a system reacts to an external or internal stimulus. When the system undergoes a step change, with transcription activated at *t* = 0, the response time is defined as the duration required for the system to reach halfway to its steady-state level. Notably, the response time (*t*_1/2_) is inversely proportional to the protein degradation rate(*32*) (see the “System response time” section in the Supplementary Materials).

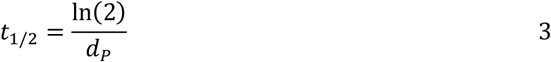

According to Equation 3, increasing *d*_*P*_ will reduce response time. In the design of the gene circuit, this parameter manipulation can be achieved by adding an aav tag to the reporter protein in the GEC (GEC-aav). An aav tag is a short peptide sequence that labels a protein as misfolded, making it more susceptible to protease-mediated degradation, thus increasing its degradation rate and reducing its half-life to ∼60 min (*35*) (Fig.1A). We simulated the dynamics of this modification using Equations 1 and 2. The predictions of this minimal model are shown in Fig.1B, where the *d*_*P*_ is increased from 0.007 to 0.07 min^−1^. As expected, the response time in the GEC-aav exhibited an apparent reduction. However, when we tested these constructs in a bulk fluorescence experiment (BFE), using superfolder yellow fluorescent protein (sfYFP) as the reporter protein, the observed dynamics deviated from the model predictions. As shown in Fig.1C, the GEC-aav unexpectedly exhibits an oscillatory behavior in a 12-hour span without external stimulations, where two peaks are observed at 230 and 550 min, respectively, and a trough is observed at 340 min. The oscillations occur with a period of 306 ± 4.96 min, estimated using autocorrelation functions (see the “Characterization of oscillations” section in the Supplementary Materials). The amplitude of the peaks reduces as time progresses, suggesting a damped oscillatory dynamic (see the “Characterization of oscillations” section in the Supplementary Materials). This unexpected oscillation suggests additional system dynamics not accounted for in the model. We observed a similar transient oscillatory profile in a separate experiment using the ssrA tag(*36*), as shown in Fig. S3, suggesting that the dynamics are unlikely to arise from tag-specific artifacts. A similar unexplained transient oscillatory response in batch culture was also reported in a classic study of negative autoregulation using a fusion protein(*37*). Although the cause was not explored by the authors, there is strong evidence that fluorescent protein fusions frequently misfold and undergo rapid degradation(*38, 39*). Together, these observations suggest that such deviations from single-scale model predictions may be more prevalent in batch culture systems than previously recognized.

**Fig. 1.**
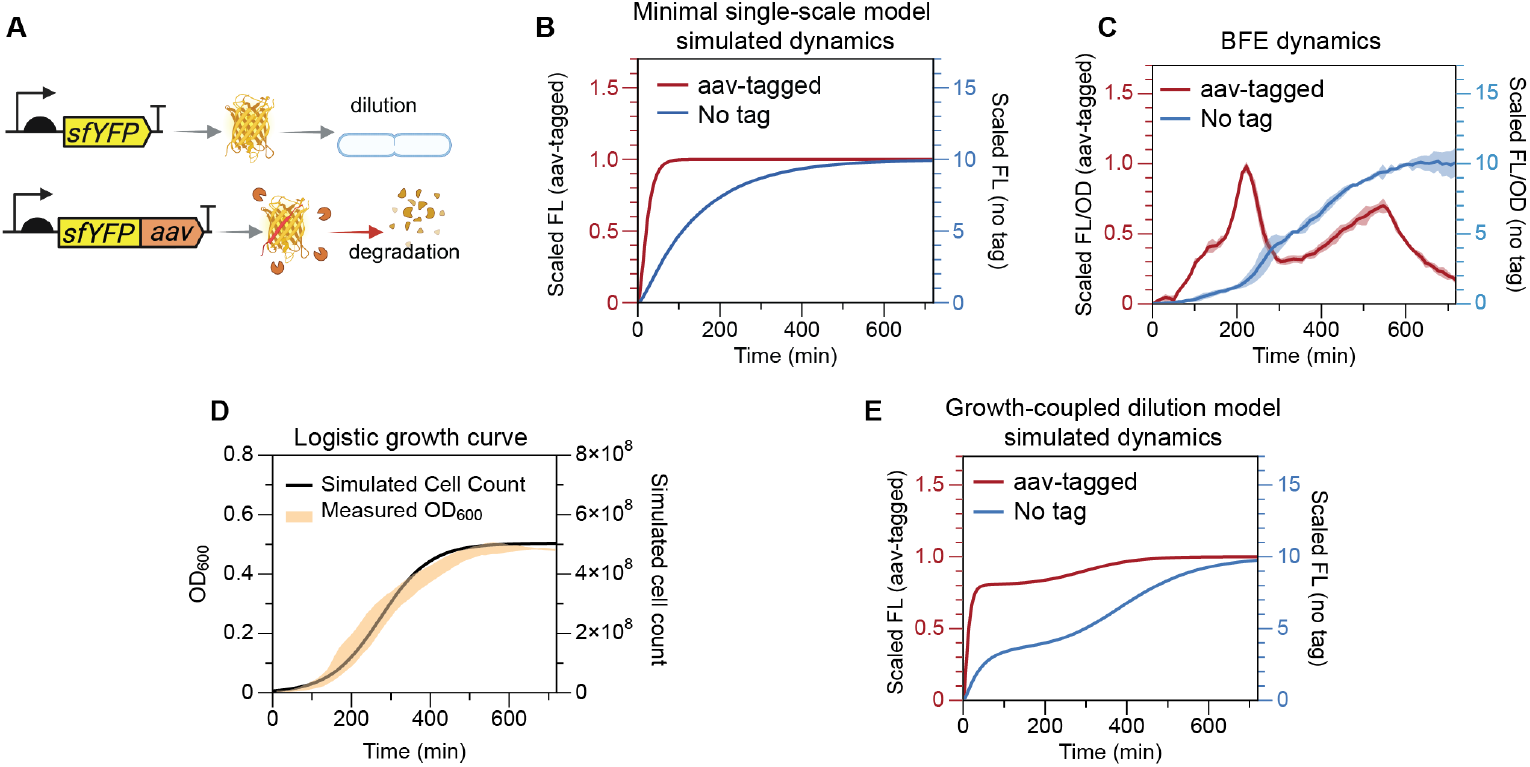
Discrepancy between experimental observations and single-scale model predictions for circuits with enhanced reporter degradation. (**A**) Circuit diagrams of the two gene expression cassettes (GECs) used in this study. For the untagged sfYFP construct, dilution due to cell growth is the predominant mode of turnover. However, for the aav-tagged sfYFP construct, both active protein degradation and dilution contribute to turnover, with degradation playing a major role compared to the untagged construct. Created in BioRender. Namboothiri, H. (2025) https://BioRender.com/y4f6l7c (**B**) Simulations using a minimal single-scale model predict faster response times as the protein degradation rate increases. (**C**) Experimentally measured fluorescence dynamics for the GEC and GEC-aav constructs reveal a clear discrepancy between model predictions and observed behavior. The GEC-aav exhibits a damped oscillatory dynamic. Solid lines show mean FL/OD, shaded bands represents standard deviation (*n* =3). (**D**) Batch culture growth, measured via optical density at 600 nm (OD_600_), compared with a logistic growth curve simulated using Equation 4. Shaded band represents one standard deviation around mean trajectory. (**E**) Simulation of a growth-coupled dilution model using Equations S7-S10 (see the “Mathematical proof showing that the growth-coupled dilution model cannot express the two-peak dynamics” section in the Supplementary Materials) fails to reproduce the experimentally observed oscillations in the aav-tagged construct. *n* denotes the number of biological replicates.

We hypothesized that this discrepancy could be due to the over-simplification of the minimal model. The model described by Equation 1 and 2 is single-scale, which only focuses on the dynamics inside a bacterial cell, with the assumption that it represents the average molecular-scale behavior of all cells in a culture that is constantly growing. The assumption of a constant growth rate, however, is not maintained in a BFE. In a BFE, cells grow in a batch culture with finite amount of nutrients and space; this results in an intricate populational dynamic that could be divided into three phases: the lag phase, the log phase, and the stationary phase. These growth dynamics can be modeled using the logistic function(*40, 41*):

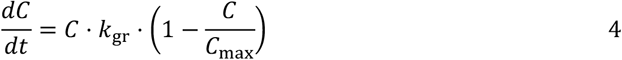

where *C* describes the cell count, *k*_*gr*_ denotes the logistic growth rate, *C*_*max*_ represents the population carrying capacity of the batch, which is limited by space and nutrient availability (Fig.1D).

Since the protein degradation/dilution rate *d*_*p*_ is dominated by cell division, it is inherently tied to the rate of cell growth. To capture this connection across scale, the overall degradation rate (*d*_*p*_) can be written as the sum of the protein degradation rate (*d*_*deg*_) and the protein dilution rate *d*_*dil*_(*C, P*), which is a function of protein concentration *P* and cell population *C*:

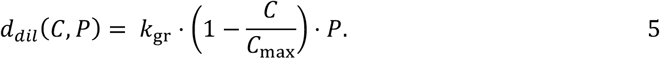

This equation relaxes the assumption that proteins are diluted at a first-order rate, instead establishing a direct connection between molecular dynamics and population-level dynamics. Similarly, mRNA dynamics also account for both intrinsic decays, *d*_*m*_, and growth-driven dilution, *d*_*dil*_(*C, M*), as described in Equation S7. By combining Equations 1 and 2 with Equation 5 and S7, we developed a growth-coupled dilution model that describes gene expression dynamics with growth-dependent dilution rate (Equation S7 to S10). As shown in Fig. 1E, the simulated dynamics of the two circuits depicted in Fig. 1A reveal a rate slow-down around *t* = 300 min in both cases. However, no oscillatory behavior was observed in the simulations.

To investigate whether the lack of oscillations using the growth-coupled dilution model is due to the parameter choice or empirical model topology, we mathematically proved by contradiction that the model is inherently monotonic, therefore cannot generate oscillatory dynamics for all parameter values (see the “Mathematical proof showing that the growth-coupled dilution model cannot express the two-peak dynamics” section in the Supplementary Materials). This finding suggests that incorporating only a growth-dependent dilution rate is insufficient for the dual-scale model to accurately capture the observed dynamics. Therefore, we hypothesize that the observed transient oscillations are emergent dynamics that rises from the growth-dependent kinetics of gene expression.

### Dual-scale CRN modeled with growth and resource-dependent rates

Growth dynamics reflects the physiological states of living cells. Therefore, in addition to dilution, they also heavily influence molecular processes such as transcription, translation, mRNA degradation, and protein degradation in the cell(*17*). We hypothesize that the transient oscillatory dynamics are driven by the multi-scale relationship between bacterial growth and growth-dependent resource allocation for gene expression. To test this hypothesis, we developed a CRN, a framework for constructing detailed mechanistic models(*42*). We propose that a dual-scale CRN, capable of capturing the intertwined dynamics of gene expression and cell growth, will successfully reproduce the transient oscillatory dynamics of sfYFP.

The dual-scale CRN integrates two physical scales: the molecular scale, which captures biochemical interactions within individual cells, and the population scale, which describes the bacterial population transitioning through growth phases (lag, exponential, and stationary). At the molecular scale, four key mechanistic steps—transcription, translation, mRNA degradation, and protein degradation—are modeled as chemical reactions, representing the average behavior across the population (see Fig. 2A-D). These steps are treated as elementary reactions governed by mass-action kinetics. At the population scale, bacterial growth is described using the logistic growth equation (Equation 4).

**Fig. 2.**
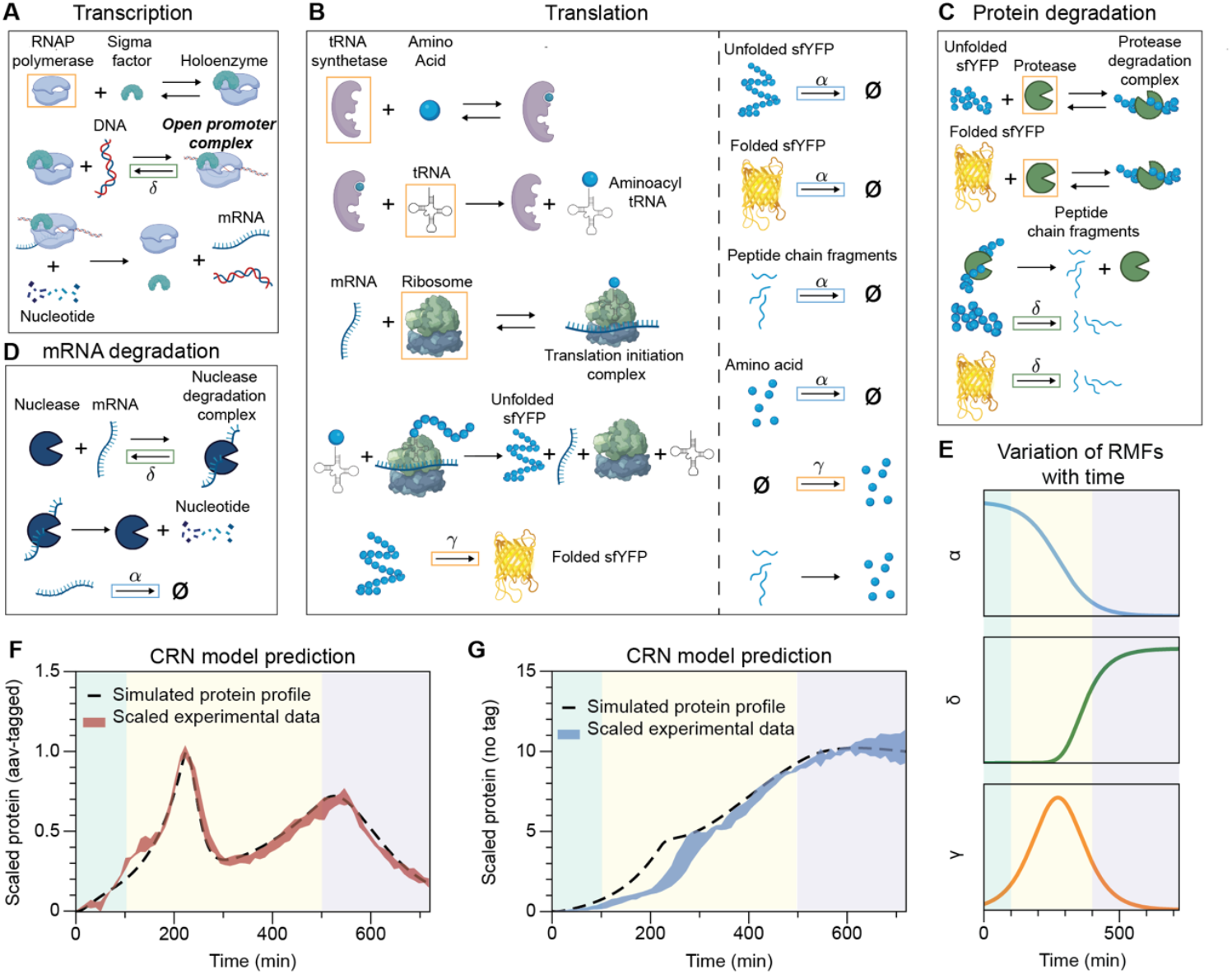
Dual-scale CRN model captures the emergent oscillatory dynamics in batch culture. (**A**–**D**) Schematic representation of the four core reaction modules in the dual-scale CRN: (**A**) transcription, (**B**) translation, (**C**) protein degradation, and (**D**) mRNA degradation. Reaction rates and molecular species modulated by RMFs are highlighted with colored boxes, corresponding to the RMFs defined in panel (**E**). Created in BioRender. Namboothiri, H. (2025) https://BioRender.com/03b2d2y (**E**) Temporal profiles of the three RMFs, α (blue), δ (green), and γ (orange), each representing a distinct growth-phase-dependent profile. The shaded background indicates the lag, log, and stationary phases. (**F**–**G**) Simulated fluorescent protein expression dynamics generated by the dual-scale CRN model (dashed lines), shown together with experimental measurements (shaded band) for GEC (**F**) and GEC-aav (**G**) (*n* =3). The model accurately reproduces the two-peak oscillatory behavior observed in the GEC-aav system and the ∼10-fold signal intensity difference between the two constructs. *n* denotes the number of biological replicates.

To couple molecular-scale dynamics with cell growth, we developed RMFs. These functions establish empirical relationships between growth-dependent steps in the CRN and the bacterial growth phases. RMFs estimate the growth phase of the population using the following equation:

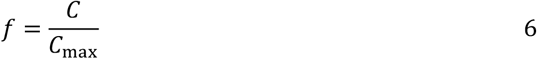

where *C* represents the current population size, and *C*_*max*_ is the population carrying capacity of the batch culture. The value of *f*, bound by 0 and 1, indicates the proximity of the population to its carrying capacity. The RMFs are functions of *f* and are used to scale the rate constants of elementary reactions, dynamically modifying rates based on the growth phase of the population. The CRN employs three distinct RMFs, which are incorporated into the four mechanistic steps as segregated in Fig. 2A-D to capture the interplay between molecular and population dynamics. In Fig. 2E we plotted the time dependent dynamics of these RMFs in a batch culture following the logistic growth function. The three sections in the figure highlights the division of growth phases, as the population transition from lag phase to log phase, then eventually to stationary phase.

The first RMF *α* has a profile described by the blue curve in Fig. 2E, defined as:

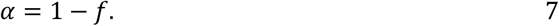

The time dependent trajectory of this RMF demonstrate a logistic growth rate of the bacterial population, where *α* portrays effective rate of cell division as the biomass accumulates in the batch culture(*4, 43*). In the CRN model, dilution of molecular species, including unfolded and folded sfYFP, mRNA, the peptide chain fragments, and the amino acids are modeled as a set of first order reactions. These reaction constants are modulated by *α*, as highlighted by the blue boxes. This coupling explicitly links molecular-scale dilution rates to population-scale dynamics in batch culture.

The second RMF *δ* is defined as:

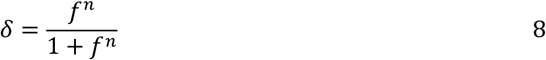

where *n* is the Hill coefficient that governs the steepness of this function (see the “Parameter estimation” section in the Supplementary Materials). Phenomenologically, *δ* represents the degree to which growth-phase transitions impact key intracellular processes, with values increasing sharply as the population approaches carrying capacity (*f* → 1). In the CRN model, *δ* modulates four reactions: In transcription (Fig. 2A), it modulates the unbinding rate of the transcription open promoter complex (OPC) into free RNAP holoenzyme and DNA, causing this step to slow down as cells transition to stationary phase, which is a well-characterized regulatory mechanism in *E. coli* for adapting to nutrient-limited environments(*44*). In mRNA degradation (Fig. 2D), *δ* modulates the unbinding rate of the nuclease degradation complex, modeling the reduced catalytic activity and increased mRNA stability observed as cells enter stationary phase(*45*). In protein degradation (Fig. 2C), *δ* modulates the degradation of both folded and unfolded proteins independent of degradation tags, broadly capturing the upregulation of endogenous proteolysis. This adjustment reflects the shift in cellular metabolism during nutrient depletion, when biosynthetic pathways are downregulated and amino acid recycling via protein degradation becomes a dominant resource strategy(*46*).

Finally, the third RMF *γ* is defined as:

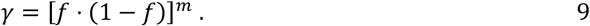

Here, *m* ∈ (0,1]. This RMF models the growth phase-dependent increase in molecular process efficiency, peaking at the highest growth rate observed during the mid-log phase. The time-dependent profile of this RMF is plotted at the bottom of Fig. 2E, where it reaches a minimum during the lag and stationary phases and peaks in the middle of the log phase, reflecting the overall scaled cell division rate as defined by the logistic growth function in Equation 4. This RMF modulates five key resource-associated species and two molecular-scale rates. The five species include RNA polymerase in transcription (*47*) (Fig. 2A), tRNA synthetase, tRNA(*20*), ribosome (*48*) in translation (Fig. 2B), and protease in protein degradation (*49*) (Fig. 2C). Notably, these five resource species are not explicitly modeled as dynamic states in the CRN; instead, the total abundance (free plus bound) is modeled as global variables that are functions of *γ*, evaluated at each time step. This represents the variation in molecular resource availability across different growth phases. Since *f* ∈ (0, 1] in the system, *f* ⋅ (1 − *f*) is bounded within the range of (0, 0.25]. If *m* = 1, then the RMF *γ* would introduce a ∼20-fold change for a modified rate as *f* transitions from 0.013 (lag phase, *γ* = 0.013) to 0.5 (log phase, *γ* = 0.25). Therefore, the exponent *m* ∈ (0, 1] adjusts the fold-change in resource abundance, allowing us to tune the dynamic range of growth phase-dependent resource allocation. Unlike the other five resources that are modified by *γ*, amino acids are explicitly modeled as a dynamic state, with their synthesis rate modulated by *γ* to reflect both growth phase-dependent effects and increased recycling due to elevated protein degradation. Finally, protein maturation rate is also modified by *γ* to capture the growth dependent fluorescent protein maturation rates in *E*.*coli(50*).

We used BioCRNpyler(*42*), a Python-based tool for CRN construction, to implement the GEAGS model in a modular and extensible format. The model and its associated RMFs were encoded in a composable structure to facilitate iterative development and refinement. In BioCRNpyler, the GEAGS framework was built by defining system components (Fig. 2A–D), specifying mechanisms that govern interactions between these components (denoted as modules in Fig. 2A– D), and assembling these into a mixture object that hosts all components and provides shared resources. To support growth phase-dependent modeling, we extended BioCRNpyler’s internal libraries by adding the additional mechanisms, introducing custom reaction propensities for the RMFs, and defining associated parameters. After parameter tuning, the resulting GEAGS model successfully reproduced the transient oscillatory dynamics observed experimentally (Fig. 2F). Using the same parameter set but setting the aav-mediated degradation rate to zero, the model also reproduced the dynamics of GEC, including both the time course and the ∼10-fold increase in signal intensity (Fig. 2G). These results demonstrate that the dual-scale GEAGS model, implemented within BioCRNpyler and incorporating three RMFs, effectively captures the interplay between growth phase-dependent resource availability, proteolysis, and dilution.

### Parameter perturbation reveals the key factors linked to transient oscillation

While the CRN model accurately captures the two-peak oscillatory dynamics, the underlying drivers of these dynamics are not immediately clear. As illustrated in Fig. 3A, the expression profile features two local maxima (points 2 and 4) and one local minimum (point 3). The first peak (point 2) is likely attributed to delayed protein accumulation due to aav-tag-mediated degradation, while the second peak (point 4) can be explained by reduced gene expression as cells transition into stationary phase. However, the dip at point 3, occurring mid-log phase, is less intuitive. In the following section, we systematically analyze all model parameters to identify the key contributors to this transient oscillatory behavior.

**Fig. 3.**
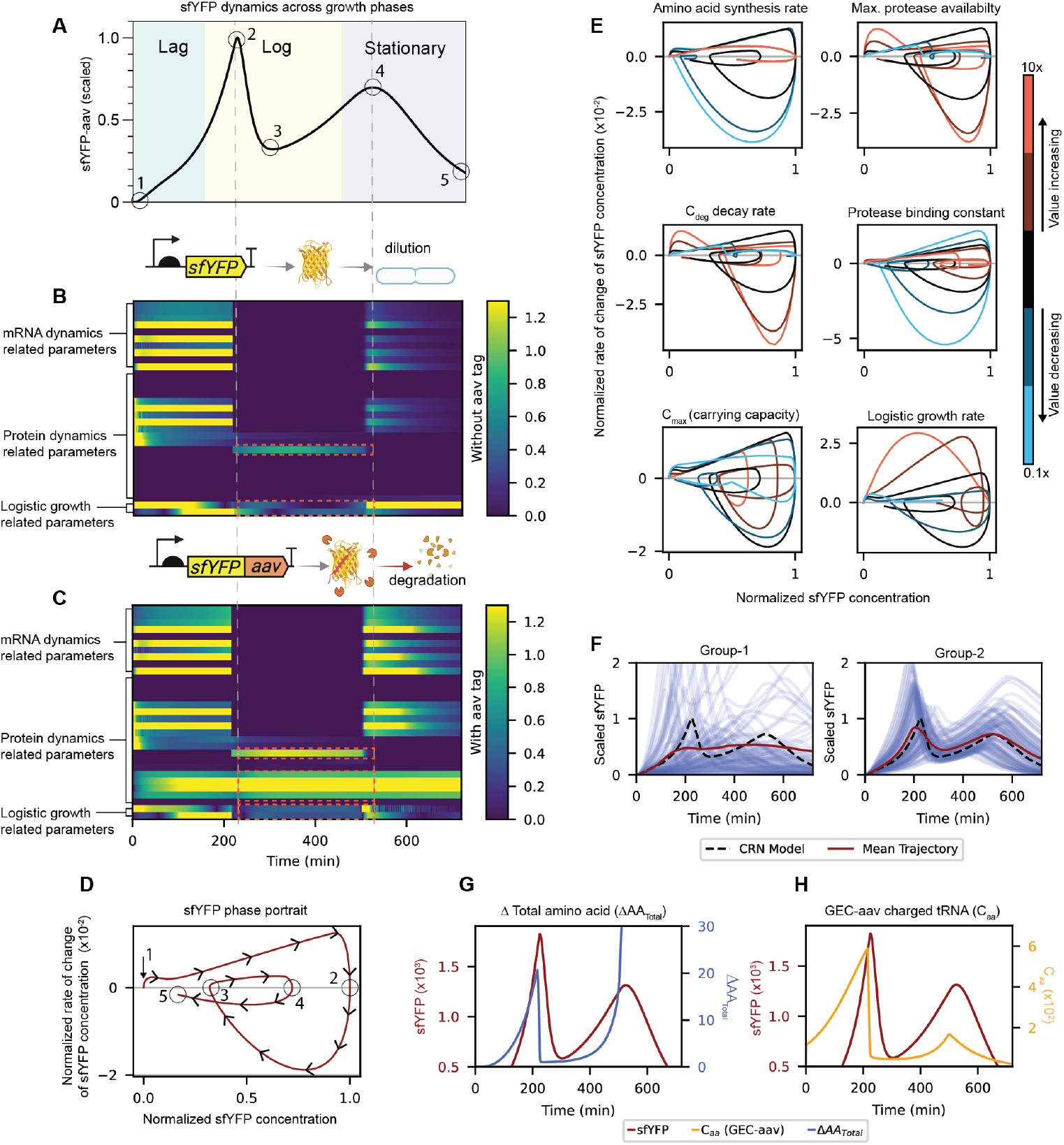
Sensitivity analysis and phase portrait diagrams reveal the mechanisms driving transient oscillation in gene expression. (**A**) Simulated GEC-aav dynamics show two peaks (points 2 and 4) and one trough (point 3) across growth phases. (**B**–**C**) Time-resolved local sensitivity analysis for untagged (GEC) and aav-tagged (GEC-aav) constructs, with parameters grouped into mRNA, protein, and growth-related categories. Seven parameters (four related to proteolytic tag, one governs amino acid synthesis, and two related to growth) show elevated sensitivity between points 2 and 4. The color scale represents the absolute local sensitivity of sfYFP dynamics with respect to the parameters. (**D**) Reporter sfYFP phase portrait illustrating oscillatory dynamics. (**E**) Phase portrait analysis (0.1× to 10× perturbation on individual parameters) reveals loss of oscillations upon perturbing *C*_*max*_, logistic growth rate, protease availability, amino acid synthesis rate, and degradation rate of the protease–protein complex. (**F**) Phase portrait analysis grouped 35 parameters by their impact on transient oscillations: Group 1 is required to preserve the two-peak phenotype, Group 2 modulates oscillatory dynamics without eliminating oscillations, and Group 3 has negligible effect. (**G**-**H**) Dynamics of translational resources supporting amino acid recycling in driving the second rise in expression. (**G**) Increased amino acid availability in GEC-aav due to protein degradation before point 3. (**H**) Charged tRNA concentration recovers before point 3.

First, we performed a local sensitivity analysis using our previously published algorithm (*51, 52*) to evaluate the time-dependent sensitivity of the reporter protein sfYFP in the GEAGS model to 35 of the 43 model parameters, leaving the parameters in RMFs fixed. The resulting sensitivity matrices for both GEC and GEC-aav variants are visualized as heatmaps (Fig. 3B, C). To improve interpretability, we grouped the parameters into three categories: mRNA dynamics-related parameters, protein dynamics-related parameters, and logistic growth-related parameters as marked on Fig. 3B, C. Our analysis focused on the time interval between points 2 and 4, as defined in Fig. 3A, which spans the oscillatory region of interest. Within this interval, the GEC-aav profile revealed seven parameters with markedly high sensitivities, suggesting that these parameters are key contributors to the oscillatory dynamics observed between the two peaks. The four protein dynamics related parameters grouped in Fig. 3C are specific to the degradation of sfYFP mediated by the aav tag. The remaining three out of the seven parameters are also highlighted in the GEC sensitivity profile in Fig. 3B. They are amino acid synthesis rate, logistic growth rate, and carrying capacity of the batch culture (*C*_max_). These three parameters play an important role in shaping the dynamics of both the constructs. While the carrying capacity and logistic growth rate determine the population-scale dynamics and thus globally influence the reaction kinetics of the system, the amino acid synthesis rate directly affects the primary resource responsible for protein synthesis, thereby exerting a direct influence on the observed dynamics.

Next, to examine the impact of broader parameter variations, we used phase portrait analysis to assess how perturbations across wide numerical ranges influence system dynamics. As shown in Fig. 3D, the sfYFP rate of change is plotted on the y-axis, and the scaled sfYFP concentration is plotted on the x-axis. In these phase portraits, the system trajectory starts at the origin and progresses clockwise, following the temporal sequence of points illustrated in Fig. 3A. The first intersection (point 1) corresponds to the beginning of the time course, where sfYFP concentrations are low. The second intersection (point 2) marks the first peak in sfYFP concentration. The third intersection (point 3) represents a trough, where the production rate begins to rise again. The fourth intersection (point 4) captures the second peak, characterized by a drop in the production rate. The trajectory terminates at point 5, indicating the endpoint of the experiment. Oscillatory behavior in the time domain corresponds to a phase portrait that intersects the x-axis three times, as seen in the representative diagram in Fig. 3D. We generated phase portraits for the key parameters identified in the sensitivity analysis, perturbing each parameter one order of magnitude (0.1×∼10× around the model parameters) as shown in Fig. 3E. Two of the parameters, the binding and unbinding rates of protease to the aav-tagged sfYFP, were combined into a single effective protease binding constant, resulting in six total phase portraits. We found that perturbations in five of these parameters disrupted the oscillatory dynamics, eliminating the local maxima and minima observed in the baseline profile. These critical parameters include the amino acid synthesis rate, the maximum protease availability, the degradation rate of the protease–protein complex, the population carrying capacity (*C*_max_), and the logistic growth rate. These findings are consistent with the local sensitivity analysis and reinforce the conclusion that these parameters are key determinants of the oscillatory behavior across a broader parameter space.

In addition to the seven key parameters, we perturbed the remaining parameters in the CRN to generate their phase portraits (Fig. S13). Based on their effects on the system’s dynamics, we categorized the parameters into three groups: 1. those that must remain within specific ranges to preserve the oscillatory dynamics, 2. those that influence the dynamics without disrupting the presence of oscillation, and 3. those that have no impact when perturbed. As a result, 18 parameters fell into the first group, where perturbation led to the loss of transient oscillation; 10 parameters belonged to the second group, affecting the dynamics but maintaining transient oscillation; and the remaining 7 parameters had no apparent impact on the dynamics (Fig. S13).

Finally, we evaluated the collective impact of simultaneously varying all parameters within each group on the system’s dynamics. This analysis explored how combined perturbations affect system behavior and highlighted the distinct roles of each parameter group(*51*). We randomly sampled each parameter from a uniform distribution bounded at ±50% of its nominal value, generating 1,000 variants per parameter. From these distributions, we randomly sampled 100 parameter sets to generate 100 unique simulation trajectories. The mean trajectory of the randomized simulations was compared with the original CRN model prediction to assess deviations (Fig. 3F). Perturbing group 1 parameters attenuated the two peaks, with the mean trajectory showing substantial deviations from the original simulation and the loss of oscillatory profile (Fig. 3F left). In contrast, perturbing group 2 parameters slightly shifted the mean trajectory but preserved the two peaks (Fig. 3F right). For group 3 parameters, no noticeable deviation from the original trajectory was observed (Fig. S14).

Taken together, our results support the hypothesis that the observed transient oscillatory dynamics arise from the interplay between growth-phase transitions and amino acid recycling driven by aav-tag-mediated protein degradation. At the first peak (point 2), protease activity catches up to sfYFP accumulation, driving the net production rate toward zero and initiating a decline in sfYFP concentration due to intensified degradation. At the trough (point 3), extensive sfYFP degradation replenishes the intracellular amino acid pool, which in turn boosts translational capacity and drives a renewed increase in protein synthesis from this local minimum. To investigate this mechanism, we examine the dynamics of two key species in Fig. 3G and Fig. 3H. In Fig. 3G, we plot the difference in total amino acid levels between the GEC-aav and GEC constructs, capturing the contribution of amino acid recycling due to aav-tag-induced protein degradation. In Fig. 3H, we show the concentration dynamics of charged tRNA (*C*_*aa*_), which is a species that directly participates in translation. In both cases, we observe that shortly before point 3, these translational resources that were initially in rapid decline undergo a turning point and begin to recover, supporting the proposed role of resource recycling in driving the upward trend after the trough. Finally, at point 4, as cells enter stationary phase, growth slows and resource limitations suppress protein synthesis, producing the second peak followed by a decline.

The multiscale dynamics of protein degradation, resource recovery, and growth-phase transitions offers a tool to interpret circuit dynamics that deviate from single-scale models. As noted earlier, a similar oscillatory signature was observed in a classic study of negative autoregulation using a fusion protein(*37*). While their system did not include an explicit degradation tag, fusion proteins are known to be more susceptible to misfolding and proteolytic degradation(*38, 39, 53*). Our results provide a mechanistic explanation for such behaviors, suggesting that growth phase-dependent resource dynamics may lead to transient oscillations when the reporter protein is subjected to proteolytic degradation. This highlights the need for dual-scale modeling frameworks when analyzing gene circuit behavior in batch culture environments.

### Modularity and adaptability of the GEAGS model framework

Although the dual-scale CRN effectively captures the oscillatory dynamics, it consists of 43 parameters and 19 equations, making it impractical for design applications. In this section, we aim to simplify the CRN into a more efficient form while establishing guidelines for constructing effective dual-scale models with RMFs.

To achieve this, we reduced the system to six equations, representing five molecular-scale species and one population-scale species for cell density (*C*). As described in Fig.4A and the “Effective GEAGS Model” section in the Supplementary Materials, the five molecular species include mRNA (*M*), sfYFP peptide chain (*P*), matured sfYFP (*P*_*m*_), degraded sfYFP in peptide chain fragments (*P*_*c*_), and amino acids (*A*). As indicated by our analysis in Fig. 3, mRNA-related parameters have a limited influence on the two-peak dynamics. Accordingly, the effective model consolidates mRNA kinetics into a single state and models protein dynamics with four states. Key molecular-scale parameters are modulated by RMFs to incorporate growth-dependent effects. We implemented three RMFs, as shown in Fig. 4C: RMF *α* (blue) modifies dilution rates of all species due to cell division and the mRNA degradation rate, capturing its decline as cells approach the stationary phase. RMF *δ* (green) modifies the protein degradation rate (*d*_*p*_), which increases as cells enter the stationary phase(*46*). RMF *γ* (orange) modifies transcription rate, translation rate, protein folding rate, aav-tag-mediated protein degradation rate, and amino acid synthesis rate. These rates are proportional to the growth rate and peak during the exponential growth phase.

**Fig. 4.**
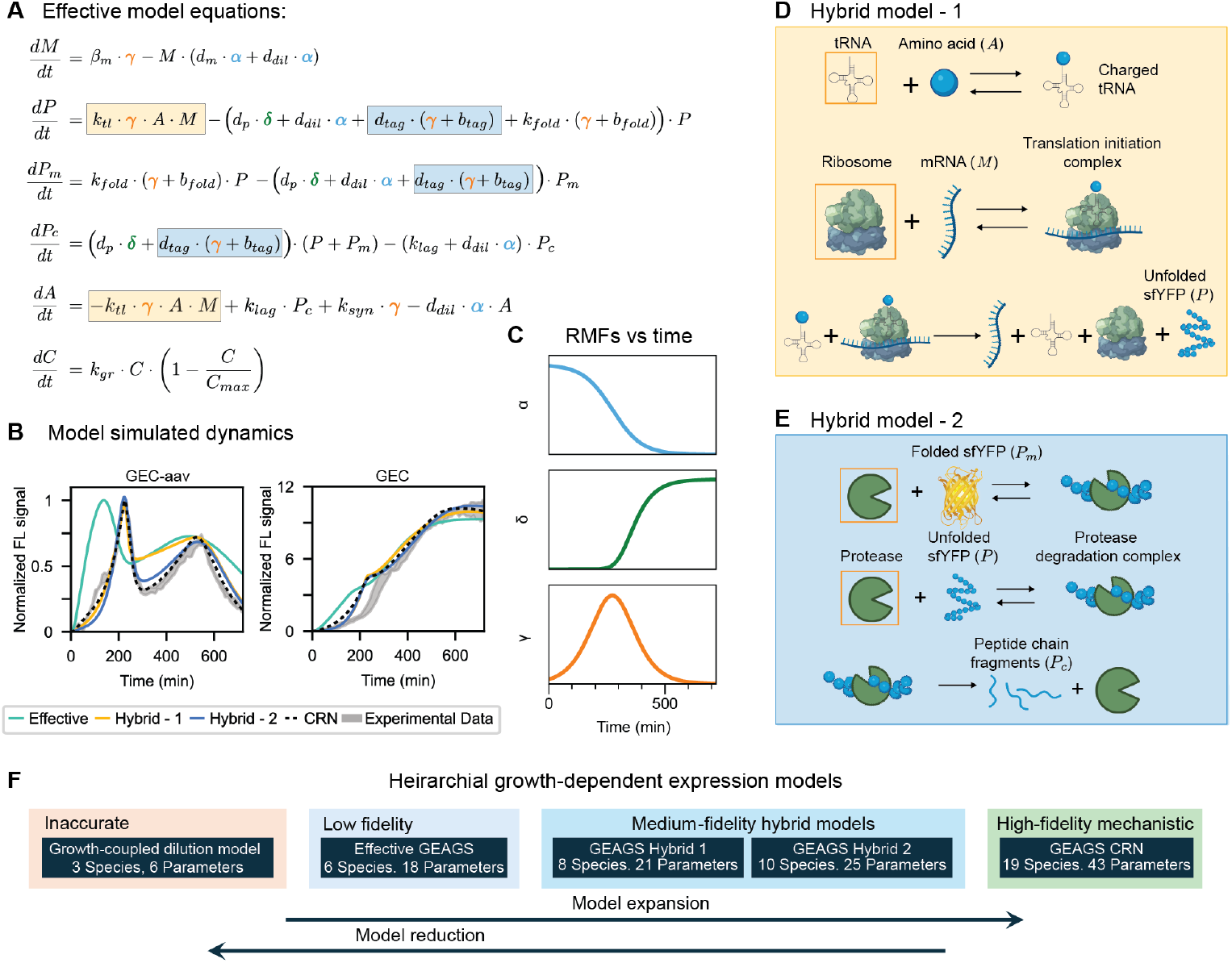
Reduction and partial-expansion of the GEAGS models. (**A**) Differential equations describing the dynamics of all species in the effective model and application of RMFs to each growth-dependent rate. Rates in boxes are later expanded in the two hybrid models. (**B**) Comparison of the effective and the two hybrid models against the CRN model simulation and experimental data. Shaded region indicates mean ± standard deviation of the experimental data (*n* = 3). (**C**) Temporal profiles of the three RMFs, *α* (blue), *δ* (green), and *γ* (orange), each representing a distinct growth-phase dependent profile. (**D**) Schematic of the expanded CRN in Hybrid model – 1. Total concentrations of the species in the orange boxes are modified by the RMF *γ*. (**E**) Schematic of the expanded CRN on protein degradation in Hybrid model – 2. Total concentrations of the species in the orange boxes are modified by the RMF *γ*. Created in BioRender. Namboothiri, H. (2025) https://BioRender.com/hcnwxgt (**F**) Hierarchy of the five GEAGS model versions developed for the GEC and GEC-aav systems in this study. *n* denotes the number of biological replicates.

Using this highly reduced effective model, we simulated the dynamics of GEC and GEC-aav, as shown in Fig. 4B (green curve). While the model qualitatively captures the oscillatory behavior and the fold-change between the two systems, it misaligns the timing of the first peak and the trough. Based on our analysis in Fig. 3, we hypothesize that this discrepancy stems from the oversimplification of translation and amino acid recycling. In the reduced model, translation is simplified as a second-order process dependent only on mRNA and amino acid concentrations. This approximation neglects key intermediate steps, including tRNA charging, ribosome binding, and elongation, leading to an overestimation of translation speed. To address this, we expanded the translation step to a CRN model that explicitly includes these intermediate processes (Fig. 4D and the “Hybrid GEAGS Model-1” section in the Supplementary Materials). In the expansion, we introduced tRNA and ribosome as part of translational resources and modeled the total concentrations of these two species as functions of *γ*. Using this hybrid model with 21 parameters and 8 equations, the simulation fit was improved (Hybrid-1 fit in Fig. 4B). The model captured the initial rise in sfYFP dynamics and correctly align the timing of the first peak. However, the timing of the trough was still misaligned (Fig. 4B, yellow). We attribute this misalignment to the oversimplified representation of aav-tag induced protein degradation and amino acid recycling, which was modeled as a modified first-order reaction in the effective model. While the RMF captures its growth-dependent nature, it likely overestimates the degradation rate. To refine this process, we further expanded Hybrid-1 model to include a two-step degradation mechanism, incorporating a reversible protease-protein binding step followed by an irreversible degradation step (Fig. 4E and the “Hybrid GEAGS Model-2” section in the Supplementary Materials). The total protease species is also modeled as a function of *γ*. With this modification (Hybrid-2), the model was expanded to 25 parameters and 10 equations. We observed that it successfully captured the trough dynamics, showing improved alignment with experimental observations and closely matching the fit achieved by the full CRN model (Fig. 4B, blue).

To assess the necessity of all three RMFs, we performed an ablation study on the effective model, removing one RMF at a time. We find that *α* and *γ* are required to reproduce the transient oscillatory dynamics, whereas the behavior persists without *δ* (Fig. S15). This is likely because *δ* models the increase in the endogenous degradation rate of sfYFP, which is extremely slow relative to the timescales of transcription, translation, dilution and degradation tag-mediated sfYFP degradation, which are modified by *α* and *γ*. Although *δ* is dispensable in this experiment, it can be important for capturing stress or burden-associated rate increases in stationary phase, reflecting nonlinear physiological adaptation.

Our results demonstrate that RMFs can be used to refine effective dual-scale models, capturing key growth-dependent dynamics with only six equations. Additionally, selectively expanding specific parameters into CRN-based representations allows for improved alignment with observed dynamics. This hybrid modeling approach provides a flexible strategy for balancing model simplicity with accuracy, ensuring better predictive performance in dual-scale systems.

### Extending the GEAGS framework to layered feedback circuits in batch culture

Finally, we sought to test whether the GEAGS framework is extensible to more complex synthetic circuits in batch culture. To this end, we applied the GEAGS framework to four constructs from our previously published study on layered feedback control (Fig. 5A)(*4*). These constructs consist of a two-stage activation cascade, in which the first gene expression cassette (GEC) encodes CinR, an activator that drives expression of sfYFP in the second GEC. The four configurations as shown in Fig.5A include: (i) an open-loop system with no feedback, (ii) a *cis* feedback system in which an antisense RNA (sRNA) (*2*) expressed from the second GEC represses its own mRNA, (iii) a *trans* feedback system in which LacI expressed from the second GEC represses the combinatorial promoter of the first GEC, and (iv) a layered feedback system that combines both motifs.

**Fig. 5.**
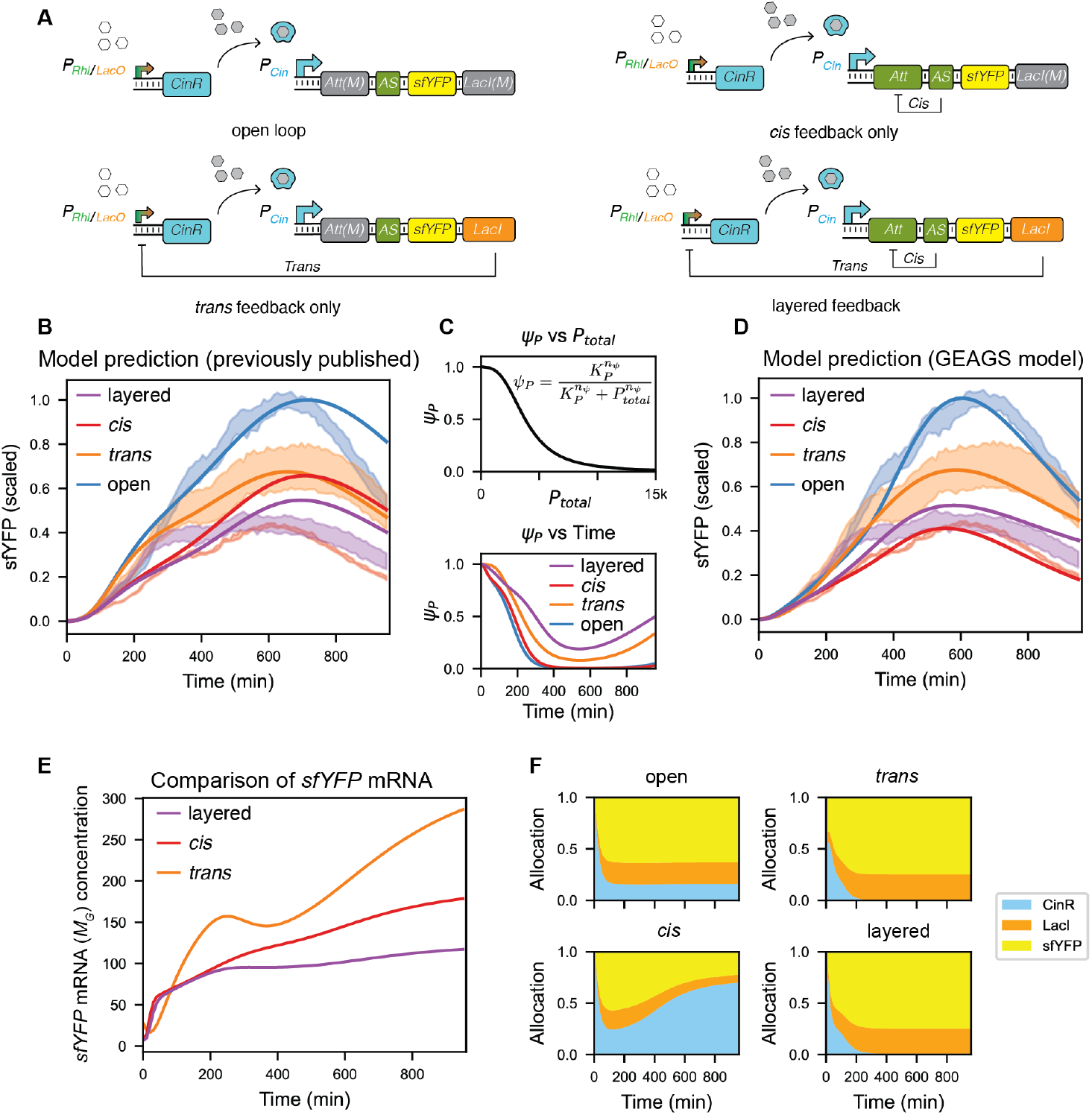
GEAGS explains resource allocation dynamics in layered feedback circuits. **(A)** Circuit diagrams for four synthetic configurations: open loop (no feedback), *trans* feedback (LacI represses CinR), *cis* feedback (sRNA represses *sfYFP* mRNA), and layered feedback (both *cis* and *trans* feedback). Figure adapted from Hu and Murray (*4*). **(B)** Predictions from previous published model fail to explain why layered feedback yields higher sfYFP expression than *cis* feedback. Data obtained from Hu and Murray (*4*). **(C)** The crowding-dependent RMF *ψ*_*p*_ decreases protein maturation and increases degradation as total recombinant protein accumulates. **(D)** The GEAGS model resolves the discrepancy, accurately reproducing sfYFP dynamics across all configurations. Data obtained from Hu and Murray (*4*). **(E)** *sfYFP* mRNA concentration is lowest in the layered feedback design compared to the two single-layered feedback systems. **(F)** Simulated translational resource allocation shows that layered feedback directs more ribosomal resources to sfYFP, while cis feedback produces more CinR, draining translational resources from sfYFP translation, yielding lower sfYFP signal.

In prior work, a single-scale model did not reproduce the observed dynamics. We then used a resource-aware model that adjusted rates by the measured growth rate (Fig. 5B). This yielded reasonable agreement for these specific circuits but was not generalizable. Notably, it left an unresolved discrepancy between the simulations and the experimental data, where the layered feedback circuit (purple shade in Fig. 5B) produced a higher sfYFP signal than the *cis*-only feedback system (red shade in Fig. 5B), contradicting both theory expectations and our earlier modeling efforts.

To revisit this system using the GEAGS framework, we constructed a 13-equation dual-scale model and applied the same three growth-dependent RMFs to capture growth-coupled dynamics in dilution, transcription, and translation rates (see the “Layered feedback control model” section in the Supplementary Materials). In addition, we introduced a growth-independent RMF, *ψ*_*P*_, to account for intracellular crowding effects that arise as recombinant protein levels accumulate over time (Fig. 5C). Specifically, *ψ*_*P*_ modulates the protein maturation rate, while the term 1 − *ψ*_*P*_ scales protein degradation, together with *δ*. As illustrated by the negative Hill function in Fig. 5C, increasing total recombinant protein concentration leads to reduced maturation efficiency, likely due to crowding-induced interference with protein folding processes(*54*). Simultaneously, crowding increases the likelihood of protein misfolding, which in turn enhances protease-mediated degradation. This mechanism explains the sharp decline in reporter signal observed in the open-loop circuit, which exhibits the highest levels of recombinant protein accumulation (blue line in Fig. 5D).

Using the GEAGS framework, we were able to resolve the discrepancy between model predictions and experimental observations for the layered feedback circuit, illustrated by the mismatch of red/purple line and red/purple shade in Fig 5B. We found that, although the layered control circuit indeed exhibits lower *sfYFP* mRNA levels than the *cis* and *trans* feedback systems (Fig. 5E), it allocated a considerably larger fraction of translational resources toward sfYFP protein synthesis (yellow region in Fig. 5F). In contrast, the *cis*-only circuit overexpressed *cinR* mRNA due to the absence of *trans* feedback, leading to competition for shared translational resources and reduced allocation to sfYFP translation. This shift in resource allocation explains why the layered control circuit ultimately exhibits higher sfYFP protein output than the cis-only system. Despite stronger transcriptional repression, elevated signal emerges due to increased allocation of translational resources to the reporter gene.

The GEAGS-based model successfully recapitulated the dynamics of all four circuit configurations (Fig. 5D), including the previously unexplained inversion where layered control resulted in higher signal than *cis*-only feedback. These results reinforce the role of growth phase-dependent resource redistribution in shaping circuit behavior and demonstrate the versatility of the GEAGS framework for modeling complex, feedback-regulated synthetic networks in non-steady-state environments.

## Discussion

The design of robust synthetic gene circuits increasingly demands modeling frameworks that capture the dynamic interplay between cellular processes and population-level physiology. In this work, we introduced GEAGS, a dual-scale modeling framework that bridges molecular gene expression and population growth dynamics via RMFs. Unlike conventional single-scale models that assume constant exponential growth, GEAGS explicitly links gene expression kinetics to batch culture growth phases, enabling accurate prediction of circuit behavior across lag, log, and stationary phases.

Our study highlights how growth phase-dependent resource redistribution can produce emergent circuit dynamics that are not captured by single-scale models. By applying GEAGS to a degradation-tagged reporter system, we resolved an unexpected oscillatory gene expression profile and identified the underlying mechanism as a combination of proteolysis-driven amino acid recycling and growth phase transition. These transient oscillations would have been obscured without integrating multiscale considerations, underscoring the power of GEAGS to reveal emergent system behaviors.

To improve accessibility for design applications, we systematically reduced the full CRN into a six-equation effective model that preserved the essential oscillatory features. We then selectively re-expanded specific modules, such as translation and degradation, to recover mechanistic fidelity where needed. This hierarchical modeling approach allows users to balance accuracy with simplicity and supports flexible design-space exploration. Moreover, we demonstrated the extensibility of GEAGS by applying it to a layered feedback control system, where the framework resolved a counterintuitive experimental outcome by uncovering how growth phase-dependent shifts in translational resource allocation reshape circuit dynamics.

This study provides a systematic recipe for building growth-dependent models of synthetic gene circuits. We recommend:(i) Start from construct-specific ODEs and include population dynamics with a logistic growth model. (ii) Scale dilution terms arising from cell growth with RMF *α*. (iii) Scale rates that rise on entry into stationary phase with RMF *δ*, for example endogenous protein degradation; if mRNA dynamics is critical, also modify the dissociation rates of open-promoter and nuclease-mRNA complexes. (iv) Apply the bell-shaped RMF *γ* to rates that peak in mid-log phase, including transcription, translation, protein maturation, amino acid biosynthesis, and degron-mediated protein degradation. In a CRN formulation, also use *γ* to scale the abundances of resource species involved in gene expression (Fig. 2). (v) To capture crowding when multiple proteins are overexpressed, scale maturation by RMF *ψ*_*P*_ and endogenous protein degradation by 1 − *ψ*_*P*_. This workflow reduces trial and error and offers a modular template for selecting and assigning RMFs to specific reactions.

These results emphasize that gene circuit behavior cannot be fully understood in isolation from the host cells’s physiological state, particularly in dynamic environments like batch cultures, where resource pools and protease activity evolve over time. GEAGS provides a modular design–build– test pipeline that integrates with tools like BioCRNpyler, making it broadly accessible to the synthetic biology community. In a separate study, we applied GEAGS to an optogenetic system in batch culture(*55*): the framework provided a priori predictions of growth-dependent dynamics and informed closed loop controller design for setpoint-tracking, and the resulting design was validated experimentally.

In its current form, however, the dual-scale coupling in GEAGS is unidirectional: molecular-scale dynamics are modulated by population-level cell growth, but not the reverse. This simplification is appropriate for systems where recombinant gene expression exerts minimal influence on host physiology. However, population dynamics are also shaped by intracellular processes(*31, 43*). This effect becomes especially prominent in circuits that actively regulate growth, especially those involving toxin–antitoxin systems (*56*) or auxotrophic rescue(*57*). Future extensions of GEAGS could incorporate this bidirectional coupling, enabling predictive modeling of synthetic circuits that modulate, as well as respond to, population growth. Such an extension would broaden the framework’s applicability to scenarios involving feedback between gene expression and population-level behavior, including population control, ecological interactions, and therapeutic containment.

Taken together, our findings establish GEAGS as a generalizable and mechanistically grounded framework that advances our ability to model, predict, and design synthetic circuits in physiologically realistic settings. As synthetic biology moves into more complex, translational domains, such multiscale modeling platforms will be essential for reliable performance and successful deployment.

## Materials and Methods

### Quantitative scaling of experimental and simulated sfYFP dynamics

For meaningful comparison between model simulations and experimental measurements, we applied a consistent scaling approach to both datasets based on the maximum fluorescence signal of the tagged sfYFP construct. Experimental fluorescence data for both GEC and GEC-aav constructs were normalized using the equation:

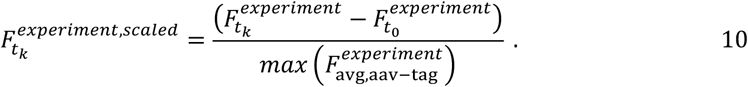

Here, 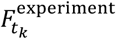 and 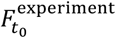 represent the optical density at 600 nm (OD_600_)-normalized fluorescence intensity at times *t*_*k*_ and *t*_0_, respectively. Subtracting the initial fluorescence value at *t*_0_ simplifies the comparison by aligning all trajectories to an assumed baseline of zero, effectively standardizing initial conditions across constructs. The denominator, 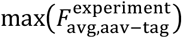 is the maximum sfYFP signal observed in the aav-tagged construct, averaged across replicates. This scaling anchors all trajectories to the GEC-aav construct’s peak signal, enabling the much higher expression level of the untagged GEC to be visualized as a relative fold increase.

Model outputs were similarly scaled using:

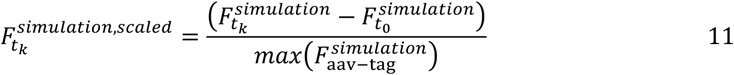

where 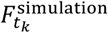 and 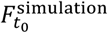 denote the simulated sfYFP concentration (in nanomolar) at times *t*_*k*_ and *t*_0_, respectively. To facilitate comparison across constructs, all trajectories were scaled relative to the maximum signal of the GEC-aav construct. This normalization enables the substantially higher expression level of the untagged GEC to be visualized on the same relative scale.

### Parameter estimation

Model parameters were estimated using a combination of manual tuning and computational optimization. The full CRN consists of 43 parameters, 21 reactions, and 19 molecular species. Among these, we measure sfYFP fluorescence, so it is the only observable state for parameter inference.

Initial parameter values were chosen based on literature sources and adjusted manually within biologically plausible bounds to qualitatively align simulated dynamics with experimental observations (see the “CRN reactions, species and parameters” section in the Supplementary Materials). Manual tuning focused on identifying parameters that most strongly influenced system behavior. Once plausible initial values for the parameters were obtained, we refined the estimates using the LMFIT(*58*), toolbox in Python. We applied a two-objective optimization strategy that simultaneously minimized two loss functions to improve the model fit.

The first loss function, *E*_1_ captures the qualitative agreement between model simulations and experimental dynamics by minimizing the sum of squared errors between scaled fluorescence trajectories:

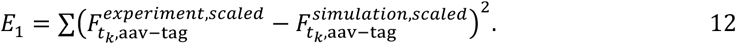

Because model simulations (in nanomolar) and experimental data (in OD-normalized fluorescence units) are not directly comparable, we introduced a second loss function, *E*_2_, to impose a quantitative constraint based on the known ∼10-fold difference in expression between the GEC and GEC-aav:

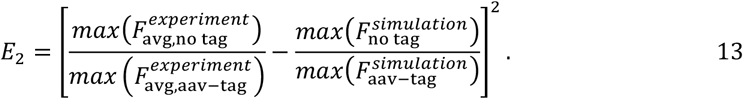

This second metric penalizes deviations in the relative expression magnitude across constructs. We defined *E*_1_ as the sum of squared errors instead of more common mean squared error to keep both the loss functions comparable, and thus, directly composable to compute a weighted loss function. The overall loss function L was defined as a weighted sum of the two metrics:

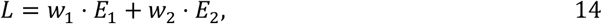

where the weights were set to *w*_1_ = *w*_2_ = 1, based on the comparable magnitudes of the two error terms across multiple trials. Optimization was performed using the Nelder–Mead (*59*) direct search algorithm, starting from manually tuned parameters. Each iteration produced updated estimates that served as the initial guess for the next round of refinement. This process was repeated until the model achieved a stable fit that align well with the experimental data.

### Phase portraits

Phase portraits were used to visualize how the rate of change of sfYFP concentration varies with respect to its concentration under large parameter spaces. Specifically, we plotted 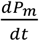 against the corresponding sfYFP concentration by simulating trajectories under varying model parameters. The time derivatives of sfYFP concentration were estimated numerically from simulation data using the gradient function in the NumPy Python library(*60*).

### Sensitivity analysis

We used local sensitivity analysis in the study to identify the key parameters most influential to the two-peak profile. We used bioscrape (*61*) to carry out the local sensitivity analysis. Bioscrape uses a previously published algorithm (*52*) in which the sensitivity coefficient matrix [*Z*_*ij*_(*t*)], is defined as:

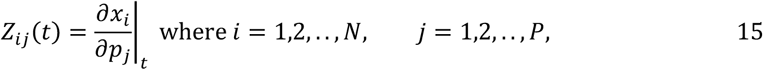

*x*_*i*_ are the model species, *p*_*j*_ are model parameters, *P* denotes the total number of parameters, and *N* denotes the total number of species. For the model, given the ODE:

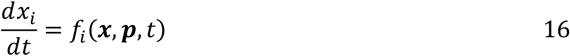

we can write *Z*_*ij*_(*t*) as the solution to the sensitivity system ODE:

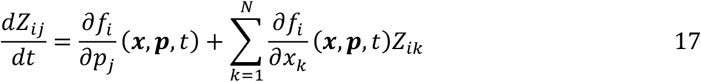

Here,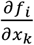 is the Jacobian matrix. We estimated the derivatives numerically using the 4^th^ order central difference method in bioscrape. Since the parameters (rate constants) in the CRN model are modified by RMFs that vary with time, to estimate the derivative 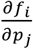 numerically, we modified the bioscrape implementation to compute sensitivity coefficients as:

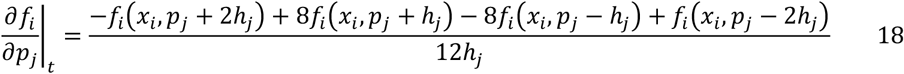

where *h*_*j*_ is computed dynamically by scaling the parameter by a small factor *h*_0_ and by the appropriate RMF for parameter *j* at the time *t* as:

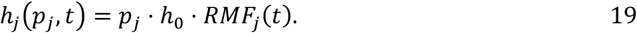

Since the only observable in the BFE is sfYFP, we focused specifically on *Z*_sfYFP,j_(*t*) to determine the key parameters that are most influential to the two-peak profile.

### Characterization of oscillations

Oscillations were analyzed using autocorrelation, discrete Fourier transform (DFT), and peak-based measurements. Autocorrelation functions were computed for each replicate and for the averaged trajectory to quantify the correlation between the signal and time-shifted versions of itself. Normalized autocorrelation was obtained using the *acf* function from the *statsmodels* library in Python. DFTs were computed for each replicate and for the averaged trajectory after mean subtraction. The DFT decomposes the signal into sinusoidal frequency components and was implemented using the *rfft* function from *scipy*.*signal* in Python. Peak-based period estimates were obtained with the *find_peaks* function from *scipy*.*signal*, and the measured period was calculated as the interval between successive peaks in the experimental traces.

### Plasmid construction and purification

All plasmids used in this study were created using Golden Gate assembly, Gibson assembly, or 3G assembly, with NEB®Turbo Competent *E. coli* as the cloning strain. Plasmids were purified using a Qiagen QIAprep Spin Miniprep Kit (Qiagen 27104).

### Strains, growth media and time-course experiments

All experiments were performed in *E. coli* strain JS006(*1*). Plasmids with different network constructs were transformed into JS006 competent cells, plated on LB + Agar plates containing 100 μg/mL carbenicillin, and incubated overnight at 37 °C. At least three colonies of each experimental condition were inoculated into 300 μL of LB containing carbenicillin in a 2 mL 96-well block and grown overnight at 37 °C at 1000 rpm in a benchtop shaker. Four microliters of the overnight culture were added to 196 μL of supplemented M9 media (1× M9 minimal salts, 1 mM thiamine hydrochloride, 0.4% glycerol, 0.2% casamino acids, 2 mM MgSO4, 0.1 mM CaCl_2_) containing carbenicillin and grown for 4 hours at 37 °C at 1000 rpm. For time course measurements, the subculture was then diluted 20x to 200 µL with M9 + carbenicillin and grown at 37 °C at 1000 rpm. During the time course, fluorescence (503 nm excitation, 540 nm emission) and OD_600_ were measured on a Biotek SynergyH1 plate reader every 10 min.

## Author contributions

Conceptualization: CYH

Methodology: CYH, AP

Investigation: HRN, CYH, AP

Visualization: HRN, CYH, AP

Supervision: CYH, AP

Writing—original draft: CYH, HRN

Writing—review & editing: CYH, HRN, AP

Resources: CYH, AP

Data curation: CYH, HRN, AP

Validation: HRN, CYH, AP

Formal analysis: HRN, CYH, AP

Funding acquisition: CYH

Project administration: CYH

## Data and materials availability

All data needed to evaluate and reproduce the conclusions in the paper are present in the paper and/or the Supplementary Materials. The source data from experiments, the source code for all the simulations, and the plasmid maps are available at the following GitHub repository: https://github.com/synbiosystems/Gene-Expression-Across-Growth-Stages and Zenodo repository: https://doi.org/10.5281/zenodo.17210634

## Supplementary materials

**This PDF file includes:**

Supplementary Text

Figs. S1 to S15

Tables S1 to S11

References (62 to 73)

