## supplemental information for "Resolving Emergent Transient Oscillations in Gene Circuits with a Growth-Coupled Model"

Hari R. Namboothiri *et al.*

* Chelsea Y. Hu.

**This PDF file includes:**

Supplementary Text
Figs. S1 to S15
Tables S1 to S11
References (62 to 73)

### S1 Characterization of Oscillations

First, we estimated the autocorrelation function of the time-series experimental data for each replicate as well as for their mean to characterize the oscillations individually. The normalized autocorrelation function was computed using the `acf` function from the *statsmodels* library in Python. The resulting autocorrelation plot (Fig. S1) indicates periodicity in the data. It is consistent with the experimentally measured periods, which were obtained by measuring the distance between successive peaks in each replicate (see Table S1). In addition, the gradual decay in the autocorrelation amplitude indicates damped oscillations, where the strength of the oscillatory behavior decreases over time.

**Table S1 Comparison of oscillation period obtained from autocorrelation functions and peak measurements.**

| | Period From Autocorrelation | Period From Peak Measurement |
|---|---|---|
| Replicate 1 | 313 min | 323 min |
| Replicate 2 | 302 min | 302 min |
| Replicate 3 | 303 min | 313 min |
| Averaged Trajectory | 313 min | 323 min |

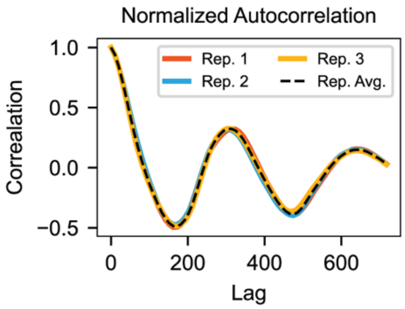


**Fig. S1.** Normalized Autocorrelation plot obtained from the data. Autocorrelation function for each replicate was estimated, by shifting the signal by lag that scales across the experimental timescale.

Next, we applied the Fast Fourier Transform (FFT) to the mean-shifted experimental data (Fig. S2A) for each replicate as well as for their mean, allowing us to characterize the oscillations individually. The analysis was conducted using the `rfft` function from the *scipy* library in Python.

The frequency spectrum from the FFT analysis (Fig. S2B) shows a clear peak at approximately 4.63e-05 Hz, corresponding to a period of 360 minutes. The reconstructed signal (Fig. S2C) demonstrates that the dominant frequency components identified by the FFT capture the oscillatory behavior of the data. Given that the time series spans only two cycles, which introduces an inherent period uncertainty of roughly 25%, this estimate is in reasonable agreement with the experimentally measured periods.

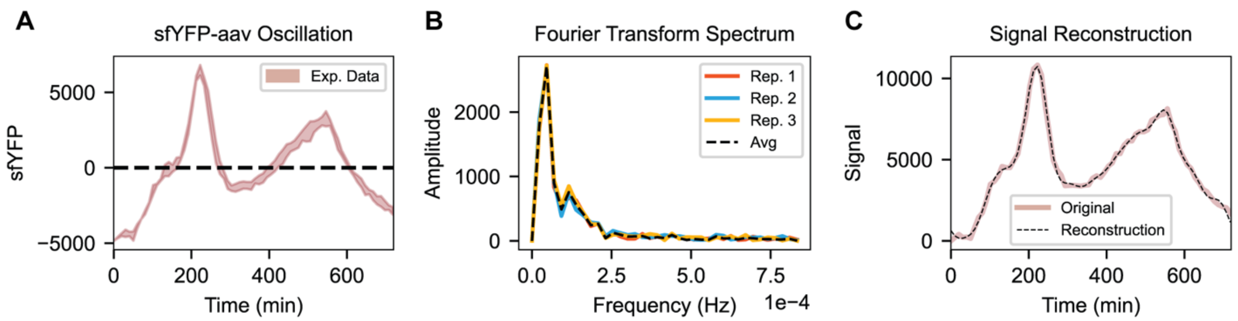


**Fig. S2.** Analysis of experimental data by Fast Fourier Transform (FFT). (A) Mean-shifted experimental data to demonstrate that the oscillations occur about the mean of the signal. (B) Spectrum of frequencies obtained by FFT. (C) Reconstruction of the original signal from one of the replicates using inverse FFT. Cosine waves generated from top 10 dominant frequencies were superimposed to obtain the reconstructed signal.

### S2 Oscillatory dynamics with ssrA tagged sfYFP

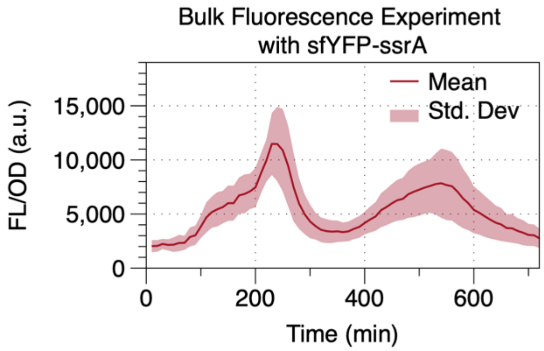


**Fig. S3.** Bulk Fluorescence Experiment with ssrA tagged sfYFP exhibits oscillatory dynamics. Solid lines show mean FL/OD, shaded bands represents standard deviation (N=4). N denotes the number of biological replicates.

### S3 Minimal Mono-Scaled and Growth-Coupled Dilution Model Analysis

#### S3.1 System Response Time

As shown in the main text, the minimal model for gene expression with two ordinary differential equations (ODEs) is:

$$\frac{dM}{dt} = \beta_m - d_m \cdot M \tag{S1}$$

$$\frac{dP}{dt} = k_{tl} \cdot M - d_p \cdot P \tag{S2}$$

Where the two species are mRNA (M) and Protein (P); P is the measurable output. $\beta_m$ and $k_{tl}$ denote the rate of transcription and translation, respectively; $d_m$ represents the degradation rate of mRNA. $d_p$ is the combined rate of degradation ($d_{deg}$) and dilution ($d_{dil}$) on protein $P$, where $d_p = d_{deg} + d_{dil}$. We assume that mRNA reaches its steady state much faster than protein does, and we can approximate equation (S2) to the following:

$$\frac{dP}{dt} = k_{tl} \cdot M_{ss} - d_p \cdot P = k_{tl} \cdot \frac{\beta_m}{d_m} - d_p \cdot P \tag{S3}$$

This ODE can be solved if we assume the initial condition of protein to 0:

$$P(t) = \frac{k_{tl} \cdot \beta_m}{d_p \cdot d_m} \cdot \left(1 - e^{-d_p \cdot t}\right) \tag{S4}$$

Where the steady state concentration of protein is:

$$P_{ss} = \frac{k_{tl} \cdot \beta_m}{d_p \cdot d_m} \tag{S5}$$

The response time of a system is the duration it takes for the system to reach halfway to its steady state. To obtain the response time for protein, we can equate equation S4 to $P_{ss}/2$ from equation S5 and solve for time:

$$P(t_{0.5}) = \frac{k_{tl} \cdot \beta_m}{d_p \cdot d_m} \cdot \left(1 - e^{-d_p \cdot t_{0.5}}\right) = \frac{P_{ss}}{2} = \frac{1}{2} \cdot \frac{k_{tl} \cdot \beta_m}{d_p \cdot d_m}$$

On rearranging the above expression we get:

$$e^{-d_p \cdot t_{0.5}} = \frac{1}{2}$$

Taking the natural logarithm on both sides, we can estimate the response time of protein to be:

$$t_{0.5} = \frac{\ln(2)}{d_p} \tag{S6}$$

Thus, the response time of protein, as predicted by the minimal model, is inversely proportional to the degradation rate of protein.

#### S3.2 Mathematical proof showing that the Growth-Coupled Dilution Model cannot express the two-peak dynamics

Consider the growth-coupled dilution model defined by the following system of ordinary differential equations (ODEs):

$$\frac{dM}{dt} = \beta - d_m \cdot M - d_{dil} \cdot M \qquad (S7)$$

$$\frac{dP}{dt} = k_{tl} \cdot M - d_p \cdot P - d_{dil} \cdot P \qquad (S8)$$

$$\frac{dC}{dt} = k_{gr} \cdot C \cdot \left(1 - \frac{C}{C_{max}}\right) \qquad (S9)$$

$$d_{dil} = k_{gr} \cdot \left(1 - \frac{C}{C_{max}}\right) \qquad (S10)$$

With initial conditions: $M(0) = P(0) = 0, C(0) = C_0$ and parameters $C_0 \in (0, C_{max})$, $\beta,\ d_m,\ k_{gr},\ k_{tl},\ d_p, C_{max} > 0$. Since the above equations describe the dynamics of a biological system, $M(t)$, $P(t)$, and $C(t)$ are continuous functions of $t$.

Goal: Empirically prove that the system's reporter protein P does not display two-peak dynamics, i.e., no pair of local maxima with a local minimum in between.

Strategy: Assume P exhibits two-peak dynamics; characterize the required critical points and curvature, then show these cannot occur under the model, thereby proving the claim by contradiction.

***Step 1: Conditions for $\boldsymbol{P(t)}$ to exhibit 2-peak dynamics:***

Let $\exists t_s = \{t_1, t_m, t_2\}$ such that $P(t)$ has two local maxima at $t_1$ and $t_2$ with $t_1 < t_2$, and a local minimum at $t_m \epsilon (t_1, t_2)$. This implies that there exists each of the following cases where:

$$\left.\frac{dP}{dt}\right|_{t=t_s} = 0\,, \qquad (S11)$$

$$\left.\frac{d^2P}{dt^2}\right|_{t=\{t_1,t_2\}} < 0 \text{ and } \left.\frac{d^2P}{dt^2}\right|_{t=\{t_m\}} > 0 \qquad (S12)$$

The second derivative of $P(t)$ can be written as:

$$\frac{d^2P}{dt^2} = k_{tl} \cdot \frac{dM}{dt} - P \cdot \frac{d(d_p + d_{dil}(t))}{dt} - (d_p + d_{dil}(t)) \cdot \frac{dP}{dt}$$

For simplicity, let:

$$\zeta(t) := \ d_p + d_{dil}(t). \qquad (S13)$$

Thus, the second derivative of $P(t)$ can be written as:

$$\frac{d^2P}{dt^2} = k_{tl} \cdot \frac{dM}{dt} - P \cdot \frac{d\zeta}{dt} - \zeta \cdot \frac{dP}{dt}. \qquad (S14)$$

At stationary points where $t = t_s$ and $\frac{dP}{dt} = 0$ :

$$\left.\frac{d^2P}{dt^2}\right|_{t=t_s} = k_{tl} \cdot \left.\frac{dM}{dt}\right|_{t=t_s} - P \cdot \left.\frac{d\zeta}{dt}\right|_{t=t_s}. \qquad (S15)$$

For Eq. S12 to be true, either $\zeta(t)$ or $M(t)$ must be non-monotonic (from Eq. S15). The monotonicity of $\zeta(t)$ and $M(t)$ will be analyzed in the next steps.

***Step 2: Analyze $\boldsymbol{\zeta(t)}$ ($\boldsymbol{\zeta(t) := d_p + d_{dil}(t)}$, from Eq. S13)***

(i) From Eq. S13 and S10, $\zeta(t) := d_p + k_{gr} \cdot \left(1 - \frac{C(t)}{C_{max}}\right)$.

(ii) The analytical solution of $C(t)$, with $C_0$ as the initial condition is:

$$C(t) = \frac{C_0 e^{k_{gr} \cdot t}}{1 + \frac{C_0}{C_{max}} \cdot \left(e^{k_{gr} \cdot t} - 1\right)}. \qquad (S16)$$

(iii) From Eq. S16, If $C_0 \in (0, C_{max})$, then $C(t) > 0$, $lim_{t \to \infty} C(t) = C_{max}$, and thus $\frac{dC}{dt} > 0 \; \forall t \geq 0$ (from Eq. S9).

(iv) From Eq. S10 and Step 2.iii, $d_{dil}(t) = k_{gr} \cdot \left(1 - \frac{C(t)}{C_{max}}\right) > 0 \; \forall t \geq 0$.

- If $d_{dil}(t)$ is a monotonic function, then one of the following conditions should be satisfied: $\left(\frac{dd_{dil}}{dt} < 0 \; \forall t \geq 0\right) \; or \; \left(\frac{d_{dil}}{dt} > 0 \; \forall t \geq 0\right)$.
- From Eq. S10:

$$\frac{dd_{dil}}{dt} = -k_{gr} \cdot \left(\frac{1}{C_{max}}\right) \cdot \frac{dC}{dt} \qquad (S17)$$

From step 2.iii: $\frac{dC}{dt} > 0 \; \forall t \geq 0$ and by definition $k_{gr}, C_{max} > 0$. This implies: $\frac{dd_{dil}}{dt} = -k_{gr} \cdot \left(\frac{1}{C_{max}}\right) \cdot \frac{dC}{dt} < 0 \;\; \forall t \geq 0$ and $lim_{t \to \infty} d_{dil}(t) = 0$.

Thus, $d_{dil}(t) \in \left(0, \; k_{gr} \cdot \left(1 - \frac{C_0}{C_{max}}\right)\right]$ is a monotonically decreasing function.

(v) $\therefore \zeta(t) := d_p + d_{dil}(t) \in \left(d_p, d_p + k_{gr} \cdot \left(1 - \frac{C_0}{C_{max}}\right)\right]$ is a monotonically decreasing function.

***Step 3: Analyze $\boldsymbol{M(t)}$***

(i) At time $t = 0$, $M(0) = 0 \implies \left.\frac{dM}{dt}\right|_{t=0} = \beta - M(0) \cdot \left(d_m + d_{dil}(0)\right) = \beta$.
$\left.\frac{dM}{dt}\right|_{t=0} = \beta > 0$. Thus, $M(t)$ increases initially.

(ii) If this continuous function is non-monotonic, it will reach a stationary point before decreasing. Assume the function reaches a point where $\left.\frac{dM}{dt}\right|_{t=t_1} = 0$.

(iii) $Let \; \exists t_1, t_2 \geq 0$ and $t_2 > t_1$ such that:

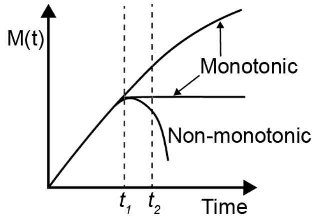


Using the above definition the following conditions can be drawn on the monotonicity of $M(t)$:

- If $M(t)$ is non-monotonic: $\exists t_2 > t_1$, such that $\left.\frac{dM}{dt}\right|_{t=t_1} = 0$, $\left.\frac{dM}{dt}\right|_{t=t_2} < 0$ , and $0 < M(t_2) < M(t_1)$.

(iv) Proof by contradiction: Assume $M(t)$ is a non-monotonic function, then show that there does not exist any $\{t_2, t_1\}$ such that $\left.\frac{dM}{dt}\right|_{t=t_1} = 0$ and $\left.\frac{dM}{dt}\right|_{t=t_2} < 0$, thus showing that $M(t)$ is a monotonically increasing function by contradiction.

From the problem statement, if $M(t)$ is a non-monotonic function, then:

$$\left.\frac{dM}{dt}\right|_{t=t_1} > \left.\frac{dM}{dt}\right|_{t=t_2} \tag{S18}$$

At time $t = t_1$,

$$\left.\frac{dM}{dt}\right|_{t=t_1} = \beta - M(t_1) \cdot \left(d_m + d_{dil}(t)\right) = 0 \tag{S19}$$

For simplicity, let $\lambda(t) \coloneqq d_m + d_{dil}(t) \in \left(d_m, d_m + k_{gr} \cdot \left(1 - \frac{C_0}{C_{max}}\right)\right]$ (From Eq. S10 and Step 2.iv). Eq. S19 can be rewritten as:

$$\left.\frac{dM}{dt}\right|_{t=t_1} = \beta - M(t_1) \cdot \lambda(t_1) = 0 \tag{S20}$$

And at time $t = t_2$,

$$\left.\frac{dM}{dt}\right|_{t=t_2} = \beta - M(t_2) \cdot \lambda(t_2) < 0 \tag{S21}$$

If $M(t)$ is a non-monotonic function, then (From Eq. S20 and S21):

$$\beta - M(t_1) \cdot \lambda(t_1) > \beta - M(t_2) \cdot \lambda(t_2)$$

$$\therefore M(t_2) \cdot \lambda(t_2) > M(t_1) \cdot \lambda(t_1) \tag{S22}$$

However, $\lambda(t)$ is a monotonically decreasing function $\left(\because \frac{d\lambda}{dt} = \frac{dd_{dil}}{dt}\ and\ \frac{dd_{dil}}{dt} < 0\ \forall t \geq 0\right)$.
$\therefore$ for $t_2 > t_1$:

$$0 < \lambda(t_2) < \lambda(t_1) \tag{S23}$$

And from problem definition:

$$0 < M(t_2) < M(t_1) \tag{S24}$$

From Eq. S23 and S24, if $0 < \lambda(t_2) < \lambda(t_1)$ and $0 < M(t_2) < M(t_1)$, then the inequality in Eq. S20 cannot be true. This contradicts the assumption that $\left.\frac{dM}{dt}\right|_{t=t_1} > \left.\frac{dM}{dt}\right|_{t=t_2}$. Hence proved by contradiction that $M(t)$ is a monotonically increasing function.

***Step 4: Proof by contradiction***

In Eq. S15:

- $\frac{dM}{dt} > 0 \; \forall t \geq 0$ (since $M(t)$ monotonically increases)
- $\frac{d\zeta}{dt} < 0 \; \forall t \geq 0$ (since $\zeta(t)$ monotonically decreases and $lim_{t\to\infty}\zeta(t) = d_p$)

Thus, the sign on $\frac{dM}{dt}$ and $\frac{d\zeta}{dt}$ will not change for $t \geq 0$. This implies that:

$$\frac{d^2P}{dt^2} > 0 \; \forall t \geq 0$$

This means that at each stationary point, the second derivative is positive, which implies that all stationary points correspond to a local minimum. This contradicts the assumption that $P(t)$ has two local maxima at $t_1$ $and$ $t_2$, with a local minimum at $t_m \in (t_1, t_2)$.

Hence, proved using contradiction that $P(t)$ cannot express two-peak dynamics.

### S4 CRN reactions, species and parameters

We divided the CRN reactions into different groups based on their mechanisms. The reactions are divided into Transcription, Translation, Amino acid replenishment, Resource maintenance, and cell growth reactions. All the reactions and rate constants are named based on the mechanism they, or the species participating in the reaction, belong to (tx, tl). They are numbered according to the order of the reactions' sequence. Subscript "b' denotes a binding step, 'u' denotes unbinding. If both are absent, then the reaction is irreversible.

#### S4.1 CRN reactions

#### S4.1.1 Transcription Reactions

$$RNAP_{free} + \sigma \rightleftarrows E_{holo} \tag{S25}$$

a. Forward rate: $r_{tx_{1b}} = k_{tx_{1b}} \cdot [RNAP_{free}] \cdot [\sigma]$
b. Reverse rate: $r_{tx_{1u}} = k_{tx_{1u}} \cdot [E_{holo}]$

The first step, binding of free RNA polymerase ($RNAP_{free}$) and sigma factor ($\sigma$), forming the RNA polymerase holoenzyme ($E_{holo}$). The binding and unbinding reaction rate constants are estimated from studies on kinetics of large protein complex formation (*62*) (protein-protein association rate of 2e6$M^{-1} \cdot s^{-1} \approx 0.12$ n$M^{-1} \cdot min^{-1}$) and holoenzyme formation (*63*) (Binding constant of ≈100-500 n$M$)

$$E_{holo} + DNA \rightleftarrows C_{open} \tag{S26}$$

a. Forward rate: $r_{tx_{2b}} = k_{tx_{2b}} \cdot [E_{holo}] \cdot [DNA]$
b. Reverse rate: $r_{tx_{2u}} = \left(k_{tx_{2u}} \cdot \left(\frac{f^n}{1+f^n}\right)\right) \cdot [C_{open}]$

The second step is binding of the holoenzyme on DNA to form the open promoter complex ($C_{open}$). The rate parameters were obtained from a study on the association rate of RNA polymerase with bacterial promoters (*64*) (reverse rate of ≈ 9.7e-2 $min^{-1}$ and dissociation constant of ≈ 100-500 nM). The parameter' $n$' determines the steepness of the change, is defined in a different context (mRNA degradation), but has been used wherever this RMF is used as a consistent measure of the steepness of change from exponential to stationary phase behavior. The estimation of n is discussed in section S3.5 of SI. Here, $f$ is a variable estimated from the logistic growth equation as:

$$f = \frac{C}{C_{max}}$$

$$C_{open} + NT \rightarrow mRNA + RNAP_{free} + \sigma + DNA \tag{S27}$$

a. Forward rate: $r_{tx_3} = k_{tx_3} \cdot [C_{open}] \cdot [NT]$

The third step is transcription elongation—an irreversible reaction. The elongation rate is estimated from the known value of 3.72kb/min (*65*) and approximated as a second-order rate constant by dividing the first-order rate by a reasonable characteristic initial nucleotide concentration (≈1000 nM).

$$mRNA + RNAase \rightleftarrows C_{deg_{mRNA}} \tag{S28}$$

a. Forward rate: $r_{tx_{4b}} = k_{tx_{4b}} \cdot [mRNA] \cdot [RNAase]$
b. Reverse rate: $r_{tx_{4u}} = \left(k_{tx_{4u}} \cdot (1+\delta)\right) [C_{deg_{mRNA}}]$

4th step is forming an mRNA degradation complex (degradosome) after binding of mRNA to RNAase enzyme complex. The determination of 'n' and the mechanism is discussed in section S3.5 of SI. The forward rate constant ($k_{tx_{4b}}$), reverse rate constant ($k_{tx_{4u}}$) and the $C_{deg_{mRNA}}$ degradation rate ($k_{tx_5}$) were estimated using guess values ($1\ min^{-1}nM^{-1}$, $100 min^{-1}$ and $1.2\ min^{-1}$ respectively) obtained from a quantitative studies on nuclease kinetics (*66*) and typical diffusion limited molecular interaction rates (*67*).

$$C_{deg_{mRNA}} \rightarrow RNAase + NT \tag{S29}$$

a. Forward rate: $r_{tx_5} = k_{tx_5} \cdot [C_{deg_{mRNA}}]$

The next step in mRNA degradation is the degradation of the degradosome into RNAase and Nucleotides. This is modeled as a first-order irreversible step.

#### S4.1.2 Translation Reactions

The first two steps in translation lead to the formation of the aminoacyl-tRNA complex ($C_{aa}$). The mechanism for the formation of $C_{aa}$ is described in a kinetics study and is also the source for the parameter estimates (*68*).

$$E_{t-free} + AA \rightleftarrows C_t \tag{S30}$$

a. Forward rate: $r_{tl_{1b}} = k_{tl_{1b}} \cdot [E_{t-free}] \cdot [AA]$
b. Reverse rate: $r_{tl_{1u}} = k_{tl_{1u}} \cdot\ [C_t]$

Binding of the amino acid ($AA$) and free enzyme aminoacyl-tRNA synthetase ($E_{t-free}$) is a reversible step.

$$C_t + tRNA_{free} \rightarrow E_{t-free} + C_{aa} \tag{S31}$$

a. Forward rate: $r_{tl_2} = k_{tl_2} \cdot [C_t] \cdot [tRNA_{free}]$

$C_t$ converts the free tRNA into $C_{aa}$ undergoing an irreversible step (*68*).

$$mRNA + Ribo_{free} \rightleftarrows C_{tic} \tag{S32}$$

a. Forward rate: $r_{tl_{3b}} = k_{tl_{3b}} \cdot [C_{aa}] \cdot [mRNA] \cdot [Ribo_{free}]$
b. Reverse rate: $r_{tl_{3u}} = k_{tl_{3u}} \cdot\ [C_{tic}]$

The next step in translation is the translation initiation. This involves forming the translation initiation complex $C_{tic}$. The parameters are estimated from a study on modeling ribosome kinetics

(*69*) and the forward rate constant is approximated similarly to reaction 3 of transcription and rates reported in typical diffusion limited molecular interactions (*67*).

$$C_{aa} + C_{tic} \rightarrow P + mRNA + Ribo_{free} + tRNA_{free} \quad \text{(S33)}$$

a. Forward rate: $r_{tl_4} = k_{tl_7} \cdot [C_{tic}] \cdot [C_{aa}]$

The translation elongation rate constant guess was estimated to be $0.8\ min^{-1}$ assuming the length of protein to be ≈300 aa and an elongation rate of 4 amino acids/s (*65*).

$$P \rightarrow P_m \quad \text{(S34)}$$

a. Forward rate: $r_{tl_5} = \left[k_{tl_5} \cdot \left(\gamma_{folding} + b_{tl_5}\right)\right] \cdot [P]$

The protein (sfYFP) folding rate was estimated from the super folder green fluorescent protein (sfGFP) maturation time of 13.6 minutes (corresponding to first order rate of $\approx 0.1\ min^{-1}$), obtained from FPbase (*70*).

Protein degradation happens in two steps first binding of the aav-tagged sfYFP (both folded and unfolded) to Protease:

$$P + P_{deg_{free}} \rightleftarrows C_{deg_P} \quad \text{(S35)}$$

$$P_m + P_{deg_{free}} \rightleftarrows C_{deg_{P_m}} \quad \text{(S36)}$$

The forward rate of binding is assumed to be different for folded and unfolded sfYFP species

a. Forward rate (unfolded sfYFP): $r_{tl_{9b_P}} = k_{tl_{9b_P}} \cdot [P] \cdot [P_{deg_{free}}]$
b. Forward rate (folded sfYFP): $r_{tl_{9b_{P_m}}} = k_{tl_{9b_{Pm}}} \cdot [P_m] \cdot [P_{deg_{free}}]$
c. Reverse rate (unfolded sfYFP): $r_{tl_{9u_P}} = k_{tl_{9u}} \cdot\ [C_{deg_P}]$
d. Reverse rate (folded sfYFP): $r_{tl_{9u_{P_m}}} = k_{tl_{9u}} \cdot\ [C_{deg_{P_m}}]$

$$C_{deg_P} \rightarrow P_c + Protease \quad \text{(S37)}$$

$$C_{deg_{P_m}} \rightarrow P_c + Protease \quad \text{(S38)}$$

a. Forward rate: $r_{tl_6} = k_{tl_6} \cdot \left[C_{deg}\right]$

$C_{deg}$ (for both folded and unfolded sfYFP) degrades into $P_c$ and Protease. $P_c$ is the species that represents degraded peptide chains that further divide into amino acids. The values for the parameters for these steps have been estimated from a study on the kinetics of the ClpXP protease and ssrA-tagged GFP (*71*).

Protein degradation independent of aav tag:

$$P \rightarrow P_c \quad \text{(S39)}$$

a. Forward rate: $r_{tl_7} = k_{tl_7} \cdot \delta \cdot [P]$ ($P$ can be both folded and unfolded protein)

This step is modeled as a first-order reaction as the protein degradation independent of aav tag need not be carried out by Protease only but could be for various reasons, so to account for that, we approximate it as a first-order step.

Degradation of peptide chain to amino acid:

$$P_c \rightarrow AA \tag{S40}$$

a. Forward rate: $r_{tl_{10}} = k_{tl_{10}} \cdot [P_c]$

The parameter for this degradation has been estimated from a kinetic study on peptidase enzyme in *E.coli (72)*.

Amino acid synthesis:

$$\phi \rightarrow AA \tag{S41}$$

a. Forward rate: $r_{tl_8} = k_{tl_8} \cdot \gamma_{syn}$

**S4.1.3 Resource Conservation Equations**

$$RNAP_{total} = RNAP_{max} \cdot \gamma_{RNAP}$$
$$RNAP_{free} = RNAP_{total} - E_{holo} \tag{S42}$$
$$E_{t\,total} = \mathrm{E}_{\mathrm{t}_{\mathrm{max}}} \cdot \gamma_{E_t}$$
$$E_{t\,free} = E_{t\,total} - C_t \tag{S43}$$
$$tRNA_{total} = \mathrm{tRNA}_{\mathrm{max}} \cdot \gamma_{\mathrm{tRNA}}$$
$$tRNA_{free} = tRNA_{total} - C_{aa} \tag{S44}$$
$$Ribo_{total} = Ribo_{max} \cdot \gamma_{Ribo}$$
$$Ribo_{free} = Ribo_{total} - C_{tic} \tag{S45}$$
$$P_{deg_{total}} = P_{deg_{max}} \cdot \gamma_{P_{deg}}$$
$$P_{deg_{free}} = P_{deg_{total}} - C_{deg_P} - C_{deg_{P_m}} \tag{S46}$$

### S4.2 Model Species

**Table S2 GEAGS CRN Model Species**

| Species | Description |
|---|---|
| $RNAP$ | RNA Polymerase |
| $\sigma$ | Transcription factor |
| $E_{holo}$ | $RNAP - \sigma$ holoenzyme |
| $DNA$ | Species representing plasmid |
| $C_{open}$ | Open promoter complex |
| $NT$ | Nucleotide species |
| $mRNA$ | Transcript for the sfYFP |
| $RNAase$ | Ribonuclease for mRNA degradation |
| $C_{deg_{mRNA}}$ | RNAase-mRNA degradation complex |
| $E_t$ | Aminoacyl tRNA synthetase |
| $AA$ | Amino acid |
| $C_t$ | $AA - E_t$ complex |
| $tRNA$ | Transfer RNA species |
| $C_{aa}$ | Aminoacyl tRNA complex |
| $Ribo$ | Ribosome species |
| $C_{tic}$ | Translation initiation complex |
| $P$ | Unfolded sfYFP |
| $P_m$ | Mature sfYFP (output signal) |
| $Protease$ | Protein degradation enzyme |
| $P_{deg}$ | Protease-Protein degradation complex |
| $P_c$ | Degraded protein (smaller peptide chain) |
| $C$ | Bacterial cell count |

### S4.3 Model Parameters

**Table S3 GEAGS CRN Model Parameters**

| Name | Description | Unit | Best fit values | Initial Estimate |
|---|---|---|---|---|
| $k_{tx_{1b}}$ | Binding rate of RNAP and Sigma factor | $1/nM \cdot min$ | 5.06e-2 | 1.2e-1 |
| $k_{tx_{1u}}$ | Unbinding rate of RNAP and Sigma factor holoenzyme | $1/min$ | 14.08 | 30 |
| $k_{tx_{2b}}$ | Binding rate of Holoenzyme and DNA | $1/nM \cdot min$ | 1.94e-4 | 3.88e-4 |
| $k_{tx_{2u}}$ | Unbinding rate of Open promoter complex | $1/min$ | 9.9-2 | 9.7-2 |
| $k_{tx_3}$ | Transcription elongation rate | $1/nM \cdot min$ | 3.6e-3 | 3.72e-3 |
| $k_{tx_{4b}}$ | Binding rate of mRNA and RNAase | $1/nM \cdot min$ | 2.55 | 1 |
| $k_{tx_{4u}}$ | Unbinding rate of mRNA-RNAase complex | $1/min$ | 161.4 | 1e2 |
| $k_{tx_5}$ | Degradation rate of mRNA-RNAase complex | $1/min$ | 0.425 | 1.2 |
| $n_\delta$ | Hill coefficient for the RMF $\delta$ | N/A | 5.5 | Estimated from data |
| $k_{tl_{1b}}$ | Binding rate of Amino acid and tRNA synthetase | $1/nM \cdot min$ | 2.79e-2 | 1e-1 |
| $k_{tl_{1u}}$ | Unbinding rate of Amino acid-tRNA synthetase complex | $1/min$ | 11.5 | 5e1 |
| $k_{tl_2}$ | Rate of formation of Aminoacylated tRNA | $1/min$ | 8.25 | 10 |
| $k_{tl_{3b}}$ | Binding rate of mRNA and Ribosome to form translation initiation complex | $1/nM \cdot min$ | 4.29e-2 | 3e-2 |
| $k_{tl_{3u}}$ | Unbinding rate of Translation initiation complex | $1/min$ | 12.34 | 5e1 |
| $k_{tl_4}$ | Translation elongation rate | $1/nM \cdot min$ | 0.17 | 8e-1 |
| $k_{tl_5}$ | Protein folding rate | $1/min$ | 0.1 | 0.07 |
| $b_{tl_5}$ | Basal Protein folding rate coefficient | N/A | 0.5 | 1 |
| $k_{tl_{9b_P}}$ | Binding rate of unfolded sfYFP and Protease | $1/nM \cdot min$ | 4.54e-4 | 1e-3 |
| $k_{tl_{9b_{P_m}}}$ | Binding rate of folded sfYFP and Protease | $1/nM \cdot min$ | 1.67e-2 | 1e-2 |
| $k_{tl_{9u}}$ | Unbinding rate of Protein-Protease complex | $1/min$ | 10.29 | 10 |
| $k_{tl_6}$ | Degradation rate of Protein-protease complex | $1/min$ | 0.39 | 1 |
| $k_{tl_7}$ | Protein degradation rate independent of aav-tag | $1/min$ | 8e-4 | 1e-3 |
| $k_{tl_{10}}$ | Peptide chain degradation to amino acid rate | $1/min$ | 1e-4 | 3e-4 |
| $k_{tl_8}$ | Amino acid synthesis rate | $nM/min$ | 138 | Empirical |
| $n_{RNAP}$ | Exponent of $\gamma$ for RNAP | $N/A$ | 0.23 | Empirical |
| $RNAP_{ma}$ | Maximum cap on RNAP concentration | $nM$ | 1.96e3 | 1e3 |
| $n_{tRNA}$ | Exponent of $\gamma$ for $tRNA$ | $N/A$ | 0.6 | Empirical |
| $tRNA_{ma}$ | Maximum cap on tRNA concentration | $nM$ | 1.535e3 | 1e3 |
| $n_{E_t}$ | Exponent of $\gamma$ for $E_t$ | $N/A$ | 0.37 | Empirical |
| $E_{t_{max}}$ | Maximum cap on $E_t$ concentration | $nM$ | 1.09e3 | 1e3 |
| $n_{Ribo}$ | Exponent of $\gamma$ for $Ribosome$ | $N/A$ | 0.67 | Empirical |
| $Ribo_{max}$ | Maximum cap on $Ribosome$ concentration | $nM$ | 280 | 1e3 |
| $n_{P_{deg}}$ | Exponent of $\gamma$ for $P_{deg}$ (Protease) | $N/A$ | 0.455 | Empirical |
| $P_{deg_{max}}$ | Maximum cap on $P_{deg}$ (Protease) concentration | $nM$ | 850 | 1e3 |
| $n_{syn}$ | Exponent of $\gamma$ for amino acid synthesis | $N/A$ | 0.296 | Empirical |
| $n_{folding}$ | Exponent of $\gamma$ for sfYFP folding | $N/A$ | 0.26 | Empirical |
| $k_{gr}$ | Logistic cell growth rate | $1/min$ | 1.578e-2 | From experiment |
| $C_{max}$ | Maximum cell growth capacity | counts | 5.037e8 | From experiment |
| $C_0$ | Initial cell count | counts | 6.67e7 | From experiment |

#### S4.4 Model simulation results with all the species

Using the above reactions and parameters, we simulated the CRN in BioCRNpyler and obtained the following results:

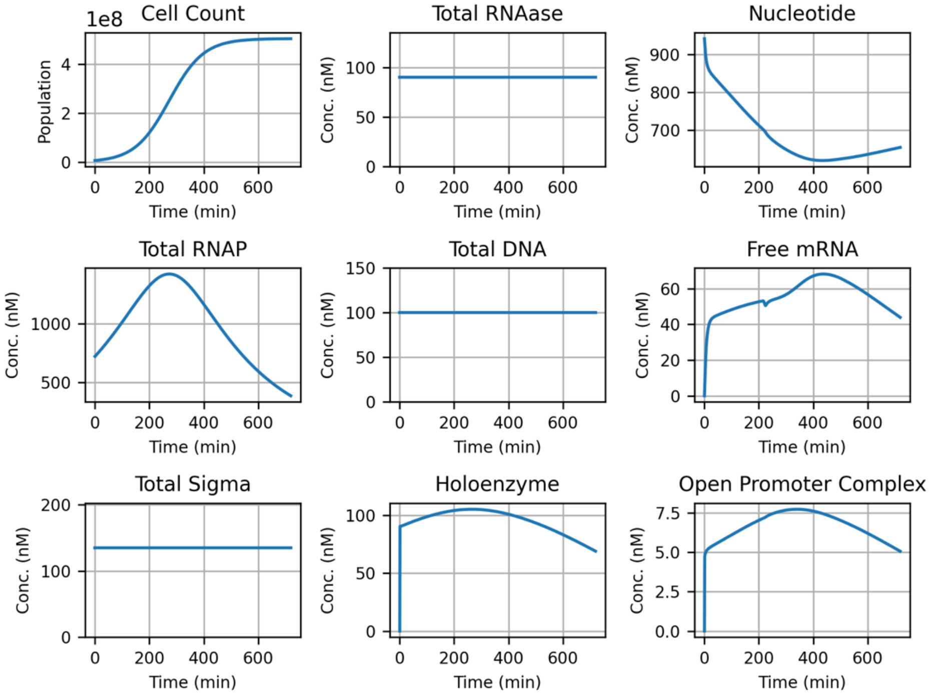


**Fig. S4.** Transcription related species in GEC

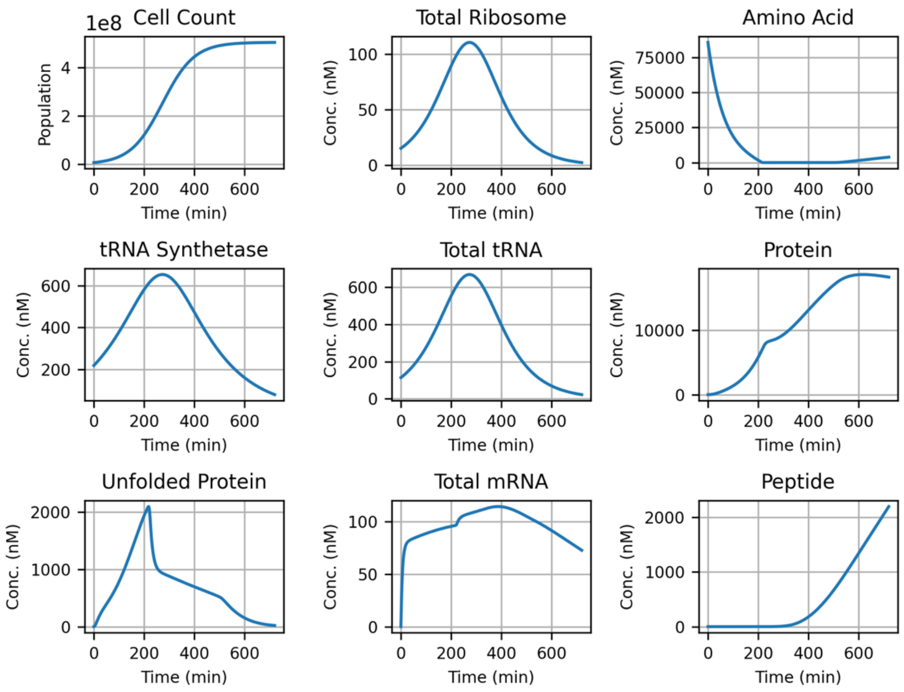


**Fig. S5.** Translation related species in GEC

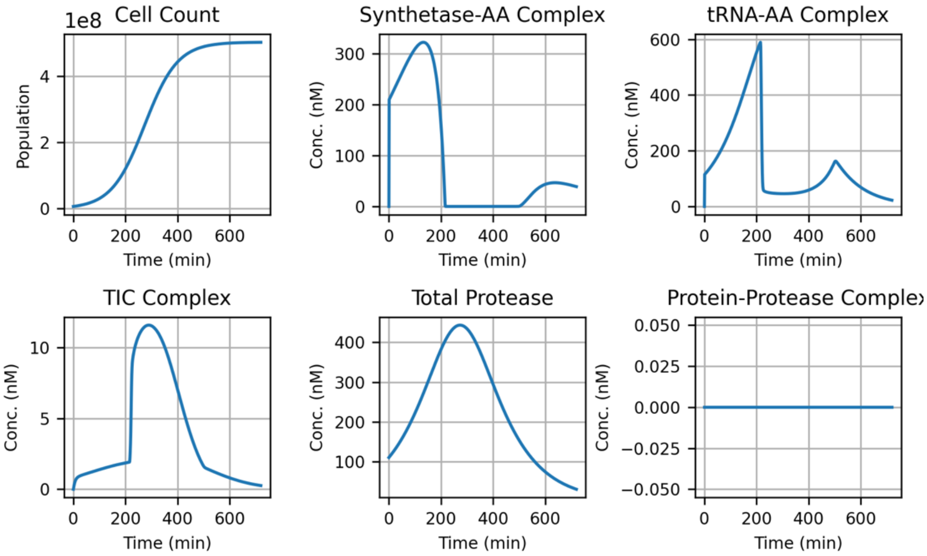


**Fig. S6.** Translation intermediate species in GEC

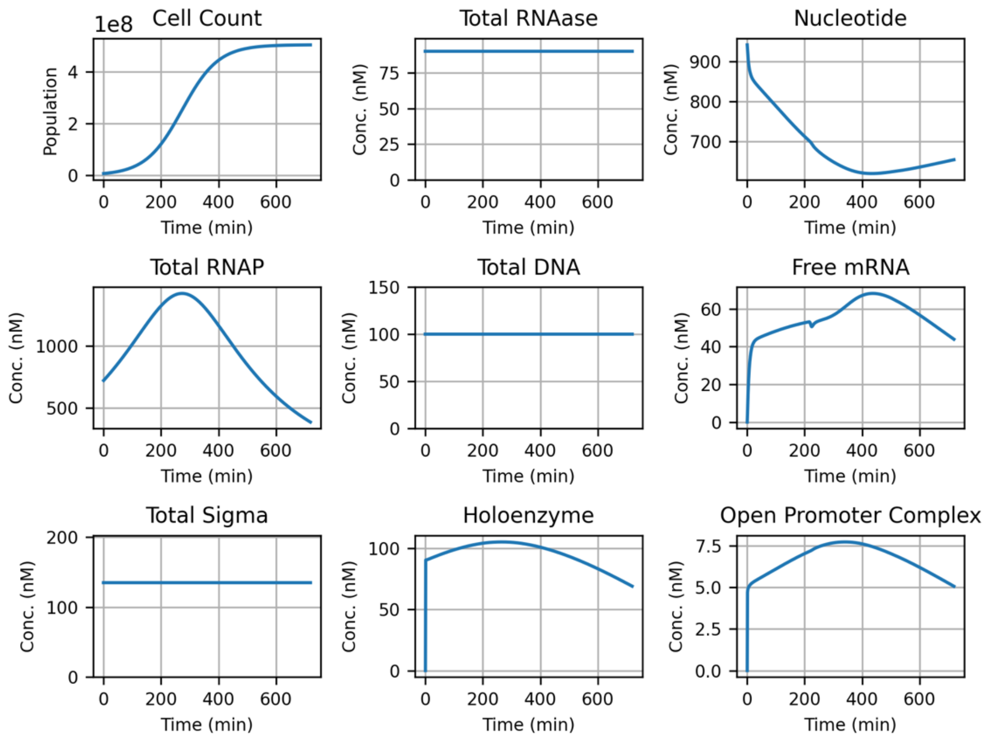


**Fig. S7.** Transcription related species in GEC-aav

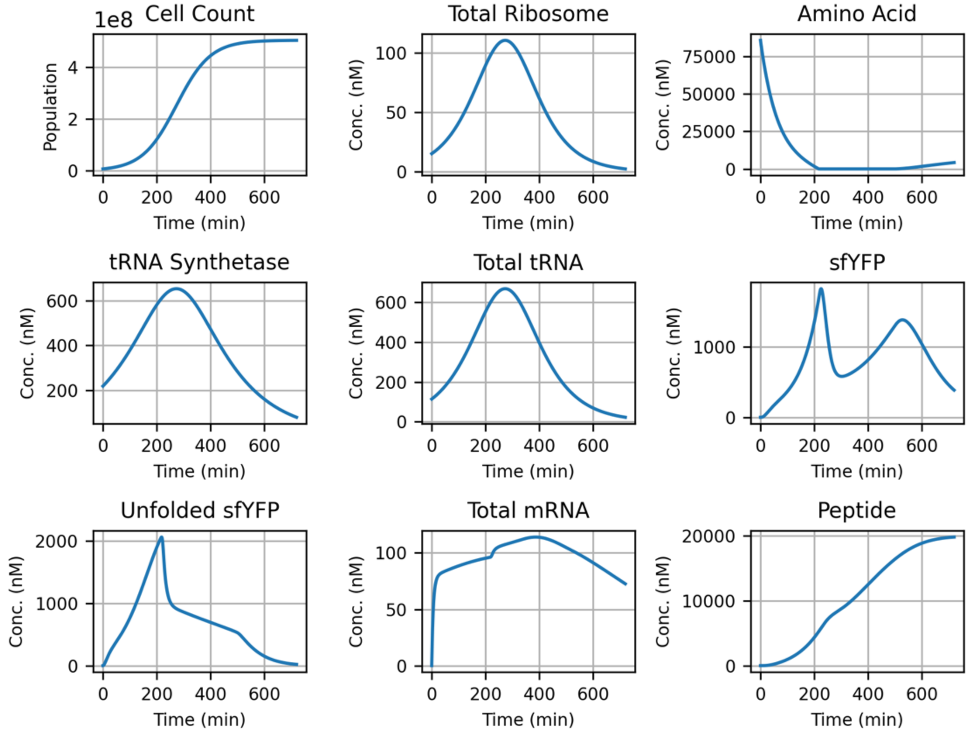


**Fig. S8.** Translation related species in GEC-aav

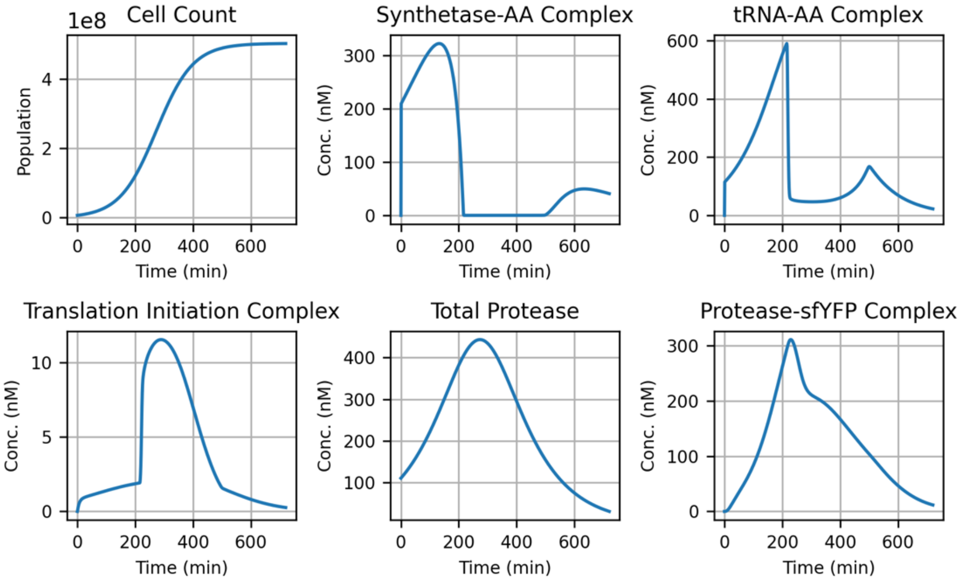


**Fig. S9.** Translation intermediate species in GEC-aav

#### S4.5 Parameter estimation

##### S4.5.1 Obtaining the Effective Range of parameters

In the CRN, many reversible reactions involve the formation of an intermediate species, whose parameters can be estimated using this method. Based on the parameter value and the concentration of the participating species, the intermediate concentration will change.To analyze this, we can consider a toy reaction:

$$A + B \rightleftharpoons C \rightarrow D + B \tag{S47}$$

Let the first reaction's forward and reverse rates be constant at $k_{+1}$ and $k_{-1}$, respectively. If we assume that the first reaction is at pseudo steady state, we can write the rate of change of C as:

$$\frac{dC}{dt} = k_{+1} \cdot [A][B] - k_{-1} \cdot [C] = 0 \tag{S48}$$

Which gives:

$$k_{+1} \cdot [A][B] = k_{-1} \cdot [C] \tag{S49}$$

We can make another assumption that as the species B is abundant and is being regenerated after the 2nd reaction, the total concentration of B remains a constant (or fixed by $\gamma$) and the amount of available B is the B that is not bound to A, which can be written as:

$$k_{+1} \cdot [A][B_{total} - C] = k_{-1} \cdot [C] \tag{S50}$$

We can rearrange the above expression to obtain the concentration of C as function of A as:

$$[C] = [B_{total}] \cdot \frac{[A]}{\frac{k_{-1}}{k_{+1}} + [A]} \tag{S51}$$

Let $\frac{k_{-1}}{k_{+1}} = K$, The variation in concentration of C with respect to A looks like:

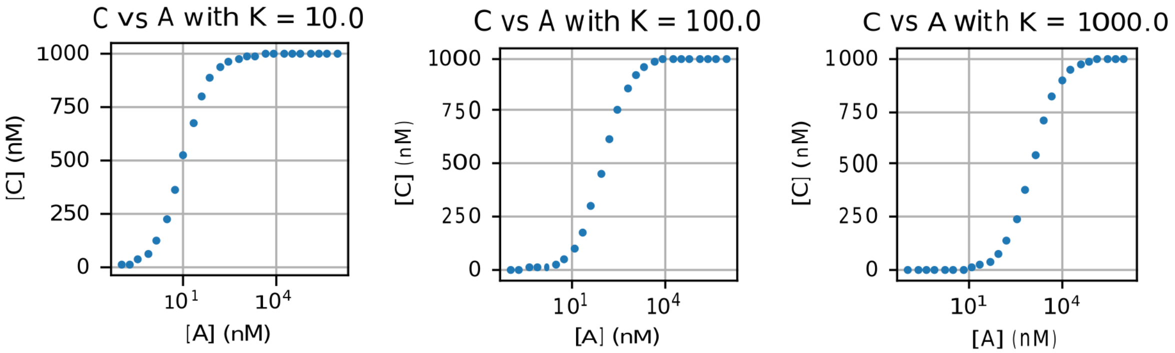


**Fig. S10.** Plot to visualize the effective range of parameters

For low or high values of [A], [C] remains unaffected, so we chose those values of parameters where for the effective range of [A], we observe the change in the [C], but also such that the parameters are within the physical range.

##### S4.5.2 Determination of the Hill coefficient for RMF $\delta$

We proposed a Hill type RMF in the reaction Eq S10 to model the decrease in the mRNA degradation with growth. The same hill-type RMF is used in Eq S8 as well. The idea of using this

RMF was derived from the data provided in a study (*73*) that measured mRNA half-lives over different growth rates (Fig S10).

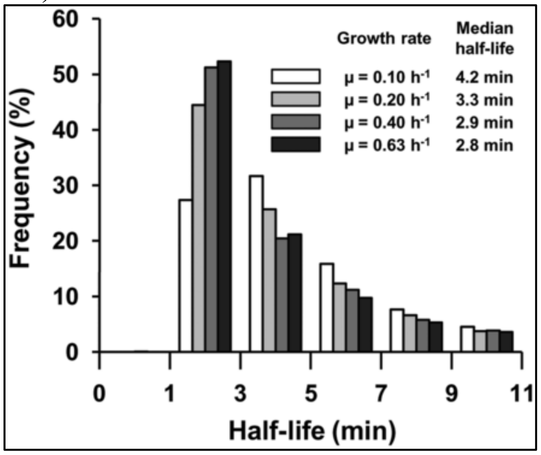


**Fig. S11.** mRNA half-lives across different growth rates. Figure taken from T.Esquerre et al. (*73*) (Fig 2)

From the half-life ($t_{0.5}$), we can estimate the first order degradation rate as:

$$d_m = \frac{\ln(2)}{t_{0.5}} \tag{S52}$$

From the logistic growth equation, we can get the specific growth rate as:

$$\mu = \frac{dC}{dt} \cdot \left(\frac{1}{C}\right) = k_{gr} \cdot \left(1 - \frac{C}{C_{max}}\right) \tag{S53}$$

We propose that the degradation rate decreases as a hill function in $f$. To model that, we use the equation:

$$d_m = \frac{d_{m_{max}}}{1 + f^n} \tag{S54}$$

Using the data and the proposed model above, we estimated the value of $n_\delta$ as 5.5 by fitting the model with the data. Fig S10 shows the agreement between the proposed model and the data.

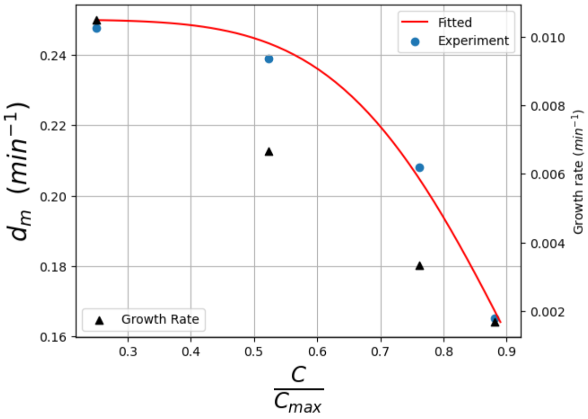


**Fig. S12.** Comparison between experimental mRNA degradation rate and model fit across different growth rates

Based on how the proposed model changes with growth, we proposed to use a similar formulation to model reactions that switch between different rates based on the growth phase they are in. We fix the parameter n to 5.5 for all the reactions to keep the dependence of the reactions on growth consistent.

### S5 CRN model analysis

#### S5.1 Phase portrait analysis plots

The parameters analyzed with phase portraits are divided into 3 groups: (1) parameters that must fall within precise ranges to maintain the double-peak dynamics (green), (2) parameters that influence the sfYFP dynamics but do not affect the existence of two-peaks (yellow), (3) parameters that do not impact the dynamics upon perturbation (violet).

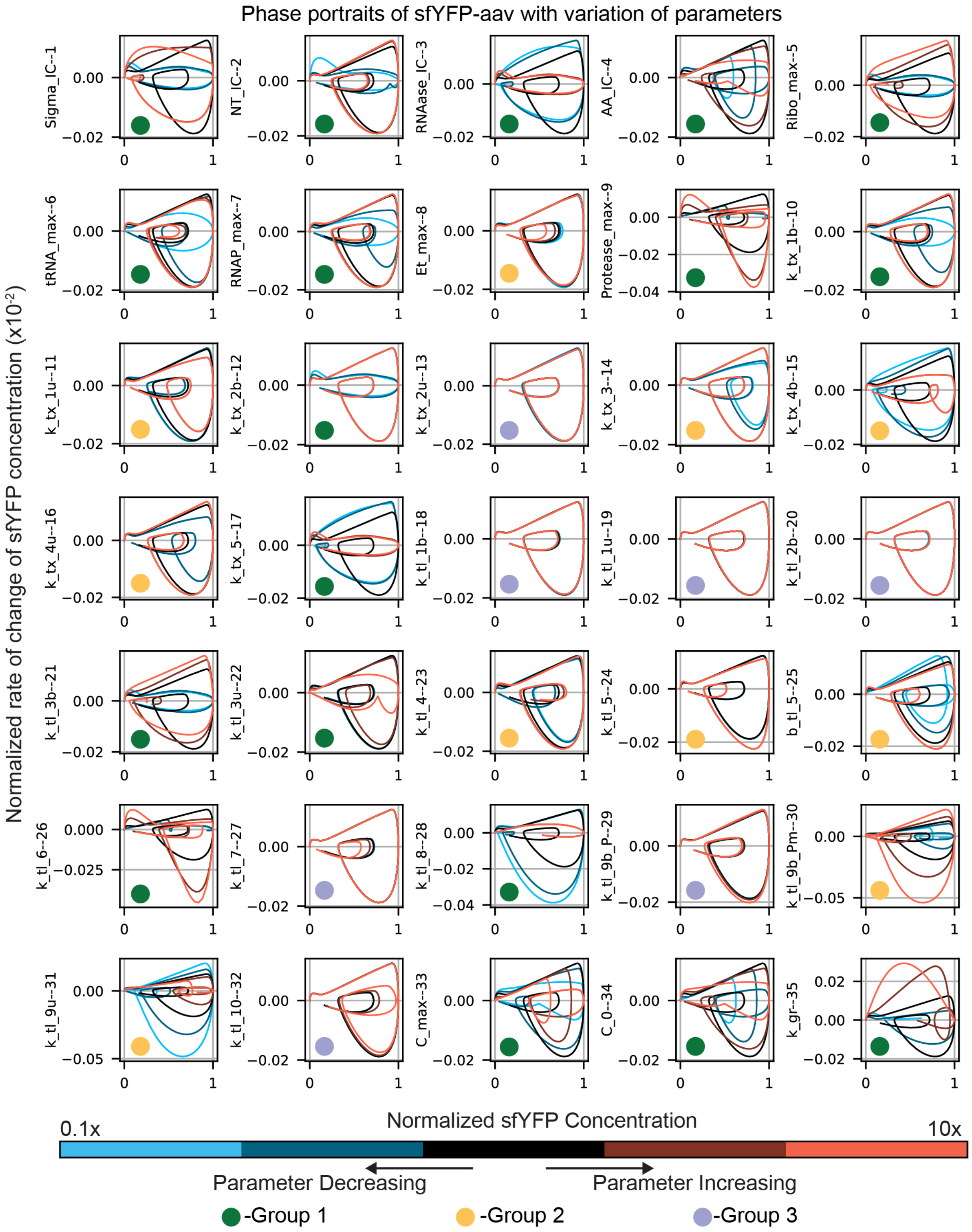


**Fig. S13.** Phase portraits obtained after varying all the parameters (0.1-10x the nominal value) in the CRN

#### S5.2 Random trajectory analysis

The 3[rd] group parameters were varied simultaneously, and the mean trajectory was plotted along with all the random trajectories. The random trajectories and the mean trajectory coincided with the original CRN trajectory indicating that these parameters do not influence the CRN dynamics.

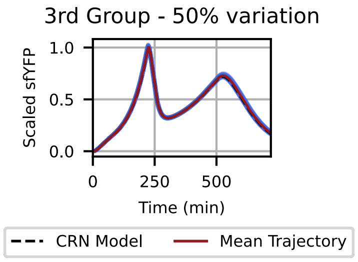


**Fig. S14.** Simulation of trajectories by varying the 3rd group parameters by 50% about the nominal value**.**

### S6 Effective GEAGS model equations and parameters

#### S6.1 Effective GEAGS Model

This section describes the minimal effective model (first model in text Figure 4). This model contains the minimal set of equations required to model the two peak dynamics. The parameters in bold indicate that they are modified by RMFs.

**Ordinary Differential Equations:**

$$\frac{dM}{dt} = \boldsymbol{\beta_m} - (\boldsymbol{d_m} + \boldsymbol{d_{dil}}) \cdot M \qquad (S55)$$

$$\frac{dP}{dt} = \boldsymbol{k_{tl}} \cdot M \cdot A - \left(\boldsymbol{d_p} + \boldsymbol{d_{tag}} + \boldsymbol{d_{dil}} + \boldsymbol{k_{fold}}\right) \cdot P \qquad (S56)$$

$$\frac{dP_m}{dt} = \boldsymbol{k_{fold}} \cdot P - \left(\boldsymbol{d_p} + \boldsymbol{d_{tag}} + \boldsymbol{d_{dil}}\right) \cdot P_m \qquad (S57)$$

$$\frac{dP_c}{dt} = \left(\boldsymbol{d_p} + \boldsymbol{d_{tag}}\right) \cdot (P + P_m) - \left(k_{l\square\square g} + \boldsymbol{d_{dil}}\right) \cdot P_c \qquad (S58)$$

$$\frac{dA}{dt} = -\boldsymbol{k_{tl}} \cdot M \cdot A + k_{lag} \cdot P_c + \boldsymbol{k_{syn}} - \boldsymbol{d_{dil}} \cdot A \qquad (S59)$$

$$\frac{dC}{dt} = k_{gr} \cdot C \cdot \left(1 - \frac{C}{C_{max}}\right) \qquad (S60)$$

**RMFs definition**

$$f = \frac{C}{C_{max}} \qquad (S61)$$

$$\alpha = 1 - f \qquad (S62)$$

$$\delta = \frac{f^{n_\delta}}{1 + f^{n_\delta}} \qquad (S63)$$

$$\gamma = \left(f \cdot (1 - f)\right)^{n_\gamma} \qquad (S64)$$

**Modified Rate Equations**

$$\boldsymbol{\beta_m} = \beta_m \cdot \gamma \quad (S65)$$
$$\boldsymbol{k_{tl}} = k_{tl} \cdot \gamma \quad (S66)$$
$$\boldsymbol{k_{fold}} = k_{fold} \cdot (\gamma + b_{fold}) \quad (S67)$$
$$\boldsymbol{k_{syn}} = k_{syn} \cdot \gamma \quad (S68)$$
$$\boldsymbol{d_m} = d_m \cdot \alpha \quad (S69)$$
$$\boldsymbol{d_p} = d_p \cdot \delta \quad (S70)$$
$$\boldsymbol{d_{tag}} = d_{tag} \cdot \gamma \quad (S71)$$
$$\boldsymbol{d_{dil}} = d_{dil} \cdot \alpha \quad (S72)$$

**Table S4 Effective GEAGS Model Species**

| Species | Description |
|---|---|
| $M$ | mRNA coding for sfYFP |
| $P$ | Unfolded sfYFP |
| $P_m$ | Folded sfYFP |
| $P_c$ | Degraded peptide chain |
| $A$ | Amino acid |
| $C$ | Cell count |

**Table S5 Effective GEAGS Model Parameters**

| Parameter | Description | Unit | Value |
|---|---|---|---|
| $\beta_m$ | Transcription rate per plasmid | $nM \cdot min^{-1}$ | 0.1 |
| $d_m$ | mRNA degradation rate constant | $min^{-1}$ | 0.16 |
| $k_{tl}$ | Translation rate | $nM^{-1} \cdot min^{-1}$ | 0.02 |
| $d_{tag}$ | Tagged protein degradation rate | $min^{-1}$ | 0.173 |
| $b_{tag}$ | Basal coefficient for $d_{tag}$ | $N/A$ | 0.055 |
| $d_p$ | Protein degradation rate independent of aav tag | $min^{-1}$ | 1e-4 |
| $k_{fold}$ | sfYFP maturation rate | $min^{-1}$ | 0.1 |
| $b_{fold}$ | Basal coefficient for $k_{fold}$ | $N/A$ | 0.5 |
| $k_{syn}$ | Amino acid biosynthesis rate | $nM.min^{-1}$ | 410 |
| $k_{lag}$ | Peptide chain fragment degradation rate | $min^{-1}$ | 8.5e-4 |
| $A_0$ | Amino acid initial condition | $nM$ | 1e5 |
| $n$ | Exponent of $\gamma$ | $N/A$ | 0.89 |
| $n_\delta$ | Hill coefficient of $\delta$ | $N/A$ | 5.5 |
| $C_0$ | Initial condition for cell population | $counts$ | 6.66e6 |
| $C_{max}$ | Max. cell population (holding capacity) | $counts$ | 5.037e8 |
| $k_{gr}$ | Logistic growth rate | $min^{-1}$ | 0.0158 |

### S6.2 Hybrid GEAGS Model – 1

This section describes the effective-CRN Hybrid model - 1 (second model in text Figure 4). In this model, we have split the translation rate into a CRN, accounting for the resource allocation for translation. The parameters in bold indicate that they are modified by RMFs.

**Ordinary Differential Equations**

$$\frac{dM}{dt} = \boldsymbol{\beta_m} - (\boldsymbol{d_m} + \boldsymbol{d_{dil}}) \cdot M - k_{tli_b} \cdot M \cdot R_{free} + k_{tli_u} \cdot C_{TIC} + k_{tl} \cdot C_{TIC} \cdot C_{aa} \qquad (S73)$$

$$\frac{dC_{aa}}{dt} = k_{aa_b} \cdot A \cdot T_{free} - k_{aa_u} \cdot C_{aa} - k_{tl} \cdot C_{TIC} \cdot C_{aa} \qquad (S74)$$

$$\frac{dC_{TIC}}{dt} = k_{tli_b} \cdot M \cdot R_{free} - k_{tli_u} \cdot C_{TIC} - k_{tl} \cdot C_{TIC} \cdot C_{aa} \qquad (S75)$$

$$\frac{dP}{dt} = k_{tl} \cdot C_{TIC} \cdot C_{aa} - \left(\boldsymbol{d_p} + \boldsymbol{d_{tag}} + \boldsymbol{d_{dil}} + \boldsymbol{k_{fold}}\right) \cdot P \qquad (S76)$$

$$\frac{dP_m}{dt} = \boldsymbol{k_{fold}} \cdot P - \left(\boldsymbol{d_p} + \boldsymbol{d_{tag}} + \boldsymbol{d_{dil}}\right) \cdot P_{\square\square} \qquad (S77)$$

$$\frac{dP_c}{dt} = \left(\boldsymbol{d_p} + \boldsymbol{d_{tag}}\right) \cdot (P + P_m) - \left(k_{lag} + \boldsymbol{d_{dil}}\right) \cdot P_c \qquad (S78)$$

$$\frac{dA}{dt} = -k_{aa_b} \cdot A \cdot T_{free} + k_{aa_u} \cdot C_{aa} + k_{lag} \cdot P_c + \boldsymbol{k_{syn}} - \boldsymbol{d_{dil}} \cdot A \qquad (S79)$$

$$\frac{dC}{dt} = k_{gr} \cdot C \cdot \left(1 - \frac{C}{C_{max}}\right) \qquad (S80)$$

**Modified Rate Equations**

$$\boldsymbol{\beta_m} = \beta_m \cdot \gamma_{rsrc} \qquad (S81)$$

$$\boldsymbol{k_{fold}} = k_{fold} \cdot \left(\gamma_{rate} + b_{fold}\right) \qquad (S82)$$

$$\boldsymbol{k_{syn}} = k_{syn} \cdot \gamma_{rate} \qquad (S83)$$

$$\boldsymbol{d_m} = d_m \cdot \alpha \qquad (S84)$$

$$\boldsymbol{d_p} = d_p \cdot \delta \qquad (S85)$$

$$\boldsymbol{d_{tag}} = d_{tag} \cdot \gamma_{rate} \qquad (S86)$$

$$\boldsymbol{d_{dil}} = d_{dil} \cdot \alpha \qquad (S87)$$

Conservation equations for growth dependent resources tRNA ($T$) and Ribosome ($R$):

$$T_{total} = T_{max} \cdot \gamma_{rsrc}$$
$$T_{free} = T_{total} - C_{aa} \qquad (S88)$$
$$R_{total} = R_{max} \cdot \gamma_{rsrc}$$
$$R_{free} = R_{total} - C_{TIC} \qquad (S89)$$

**Table S6 Hybrid GEAGS Model - 1 Species**

| Species | Description |
|---|---|
| $M$ | mRNA coding for sfYFP |
| $T$ | tRNA species |
| $R$ | Ribosome species |
| $P$ | Unfolded sfYFP |

| | |
|---|---|
| $P_m$ | Folded sfYFP |
| $P_c$ | Degraded peptide chain |
| $A$ | Amino acid |
| $C$ | Cell count |

**Table S7 Hybrid GEAGS Model - 1 Parameters**

| Parameter | Description | Unit | Value |
|---|---|---|---|
| $\beta_m$ | Transcription rate per plasmid | $nM \cdot min^{-1}$ | 2.6 |
| $d_m$ | mRNA degradation rate constant | $min^{-1}$ | 0.1 |
| $k_{aa_b}$ | tRNA and Amino acid binding rate | $nM^{-1} \cdot min^{-1}$ | 1 |
| $k_{aa_u}$ | Charged tRNA unbinding rate | $min^{-1}$ | 100 |
| $k_{tli_b}$ | Ribosome and mRNA binding rate | $nM^{-1} \cdot min^{-1}$ | 0.07 |
| $k_{tli_u}$ | Ribosome and mRNA unbinding rate | $min^{-1}$ | 600 |
| $k_{tl}$ | Translation elongation rate | $nM^{-1} \cdot min^{-1}$ | 0.1 |
| $d_{tag}$ | Tagged protein degradation rate | $min^{-1}$ | 0.14 |
| $b_{tag}$ | Basal coefficient for $d_{tag}$ | $N/A$ | 0.1 |
| $d_p$ | Protein degradation rate independent of aav tag | $min^{-1}$ | 8e-4 |
| $k_{fold}$ | sfYFP maturation rate | $min^{-1}$ | 0.27 |
| $b_{fold}$ | Basal coefficient for $k_{fold}$ | $N/A$ | 1 |
| $k_{syn}$ | Amino acid biosynthesis rate | $nM.min^{-1}$ | 200 |
| $k_{lag}$ | Peptide chain fragment degradation rate | $min^{-1}$ | 1.6e-3 |
| $A_0$ | Amino acid initial condition | $nM$ | 8e4 |
| $n_{rsrc}$ | Exponent of $\gamma$ for resources | $N/A$ | 0.5 |
| $n_{rate}$ | Exponent of $\gamma$ for rates | $N/A$ | 0.5 |
| $n_\delta$ | Hill coefficient of $\delta$ | $N/A$ | 5.5 |
| $C_0$ | Initial condition for cell population | $counts$ | 6.66e6 |
| $C_{max}$ | Max. cell population (holding capacity) | $counts$ | 5.037e8 |
| $k_{gr}$ | Logistic growth rate | $min^{-1}$ | 0.0158 |
| $tRNA_{max}$ | Max. total tRNA availability | $nM$ | 600 |
| $Ribo_{max}$ | Max. total Ribosome availability | $nM$ | 150 |

#### S6.3 Hybrid GEAGS Model – 2

This section describes the effective-CRN Hybrid Model - 2 (third model in Fig. 4). This model is an extension of the Hybrid Model – 1, where we further split the protein degradation rate into a CRN that accounts for the growth dependent availability of Protease and the subsequent binding-unbinding steps with the tagged protein.

**Ordinary Differential Equations**

$$\frac{dM}{dt} = \boldsymbol{\beta_m} - (\boldsymbol{d_m} + \boldsymbol{d_{dil}}) \cdot M - k_{tli_b} \cdot M \cdot R_{free} + k_{tli_u} \cdot C_{TIC} + k_{tl} \cdot C_{TIC} \cdot C_{aa} \quad (S90)$$

$$\frac{dC_{aa}}{dt} = k_{aa_b} \cdot A \cdot T_{free} - k_{aa_u} \cdot C_{aa} - k_{tl} \cdot C_{TIC} \cdot C_{aa} \quad (S91)$$

$$\frac{dC_{TIC}}{dt} = k_{tli_b} \cdot M \cdot R_{free} - k_{tli_u} \cdot C_{TIC} - k_{tl} \cdot C_{TIC} \cdot C_{aa} \quad (S92)$$

$$\frac{dP}{dt} = k_{tl} \cdot C_{TIC} \cdot C_{aa} - \left(\boldsymbol{d_p} + \boldsymbol{d_{tag}} + \boldsymbol{d_{dil}} + \boldsymbol{k_{fold}}\right) \cdot P \quad (S93)$$

$$\frac{dP_m}{dt} = \boldsymbol{k_{fold}} \cdot P - \left(\boldsymbol{d_p} + \boldsymbol{d_{tag}} + \boldsymbol{d_{dil}}\right) \cdot P_m \quad (S94)$$

$$\frac{dC_{deg_{P_m}}}{dt} = k_{deg_{bP_m}} \cdot P_m \cdot P_{deg_{free}} - k_{deg_u} \cdot C_{deg_{P_m}} - k_{c_{deg}} \cdot C_{deg_{P_m}} \quad (S95)$$

$$\frac{dC_{deg_P}}{dt} = k_{deg_{bP}} \cdot P \cdot P_{deg_{free}} - k_{deg_u} \cdot C_{deg_P} - k_{c_{deg}} \cdot C_{deg_P} \quad (S96)$$

$$\frac{dP_c}{dt} = \boldsymbol{d_p} \cdot (P + P_m) + k_{c_{deg}} \cdot \left(C_{deg_{P_m}} + C_{deg_P}\right) - \left(k_{lag} + \boldsymbol{d_{dil}}\right) \cdot P_c \quad (S97)$$

$$\frac{dA}{dt} = -k_{aa_b} \cdot A \cdot T_{free} + k_{aa_u} \cdot C_{aa} + k_{lag} \cdot P_c + \boldsymbol{k_{syn}} - \boldsymbol{d_{dil}} \cdot A \quad (S98)$$

$$\frac{dC}{dt} = k_{gr} \cdot C \cdot \left(1 - \frac{C}{C_{max}}\right) \quad (S99)$$

**Modified Rate Equations**

$$\boldsymbol{\beta_m} = \beta_m \cdot \gamma_{rsrc} \quad (S100)$$

$$\boldsymbol{k_{fold}} = k_{fold} \cdot \left(\gamma_{rate} + b_{fold}\right) \quad (S101)$$

$$\boldsymbol{k_{syn}} = k_{syn} \cdot \gamma_{rate} \quad (S102)$$

$$\boldsymbol{d_m} = d_m \cdot \alpha \quad (S103)$$

$$\boldsymbol{d_p} = d_p \cdot \delta \quad (S104)$$

$$\boldsymbol{d_{dil}} = d_{dil} \cdot \alpha \quad (S105)$$

Conservation equations for growth dependent resources tRNA ($T$) and Ribosome ($R$):

$$T_{total} = T_{max} \cdot \gamma_{rsrc}$$
$$T_{free} = T_{total} - C_{aa} \quad (S106)$$
$$R_{total} = R_{max} \cdot \gamma_{rsrc}$$
$$R_{free} = R_{total} - C_{TIC} \quad (S107)$$
$$P_{deg_{total}} = P_{dge} \cdot \gamma_{deg}$$
$$P_{deg_{free}} = P_{deg_{total}} - C_{deg_P} - C_{deg_{P_m}} \quad (S108)$$

**Table S8 Hybrid GEAGS Model - 2 Species**

| Species | Description |
|---|---|
| $M$ | mRNA coding for sfYFP |
| $T$ | tRNA species |
| $R$ | Ribosome species |
| $P_{deg}$ | Protease species |
| $P$ | Unfolded sfYFP |
| $P_m$ | Folded sfYFP |
| $P_c$ | Degraded peptide chain |
| $A$ | Amino acid |
| $C$ | Cell count |

**Table S9 Hybrid GEAGS Model - 2 Parameters**

| Parameter | Description | Unit | Value |
|---|---|---|---|
| $\beta_m$ | Transcription rate per plasmid | $nM \cdot min^{-1}$ | 2.6 |
| $d_m$ | mRNA degradation rate constant | $min^{-1}$ | 0.17 |
| $k_{aa_b}$ | tRNA and Amino acid binding rate | $nM^{-1} \cdot min^{-1}$ | 0.05 |
| $k_{aa_u}$ | Charged tRNA unbinding rate | $min^{-1}$ | 100 |
| $k_{tli_b}$ | Ribosome and mRNA binding rate | $nM^{-1} \cdot min^{-1}$ | 0.03 |
| $k_{tli_u}$ | Ribosome and mRNA unbinding rate | $min^{-1}$ | 600 |
| $k_{tl}$ | Translation elongation rate | $nM^{-1} \cdot min^{-1}$ | 0.17 |
| $d_p$ | Protein degradation rate independent of aav tag | $min^{-1}$ | 8e-4 |
| $k_{fold}$ | sfYFP maturation rate | $min^{-1}$ | 0.16 |
| $b_{fold}$ | Basal coefficient for $k_{fold}$ | $N/A$ | 0.7 |
| $k_{syn}$ | Amino acid biosynthesis rate | $nM.min^{-1}$ | 110 |
| $k_{deg_{bP_m}}$ | Protease and $P_m$ binding rate | $nM^{-1} \cdot min^{-1}$ | 7e-4 |
| $k_{deg_{bP}}$ | Protease and $P$ binding rate | $nM^{-1} \cdot min^{-1}$ | 1e-5 |
| $k_{deg_u}$ | Protease degradation complex unbinding rate | $min^{-1}$ | 0.45 |
| $k_{c_{deg}}$ | Protease degradation complex degradation rate | $min^{-1}$ | 0.37 |
| $k_{lag}$ | Peptide chain fragment degradation rate | $min^{-1}$ | 9.9e-4 |
| $A_0$ | Amino acid initial condition | $nM$ | 9.5e4 |
| $n_{rsrc}$ | Exponent of $\gamma$ for resources | $N/A$ | 0.56 |
| $n_{rate}$ | Exponent of $\gamma$ for rates | $N/A$ | 0.2 |
| $n_{deg}$ | Exponent of $\gamma$ for degradation | $N/A$ | 0.34 |
| $n_\delta$ | Hill coefficient of $\delta$ | $N/A$ | 5.5 |
| $C_0$ | Initial condition for cell population | $counts$ | 6.66e6 |
| $C_{max}$ | Max. cell population (holding capacity) | $counts$ | 5.037e8 |
| $k_{gr}$ | Logistic growth rate | $min^{-1}$ | 0.0158 |
| $tRNA_{max}$ | Max. total tRNA availability | $nM$ | 980 |
| $Ribo_{max}$ | Max. total Ribosome availability | $nM$ | 350 |
| $P_{deg_{max}}$ | Max. total Protease availability | $nM$ | 980 |

### S7 Ablation study on reduced models

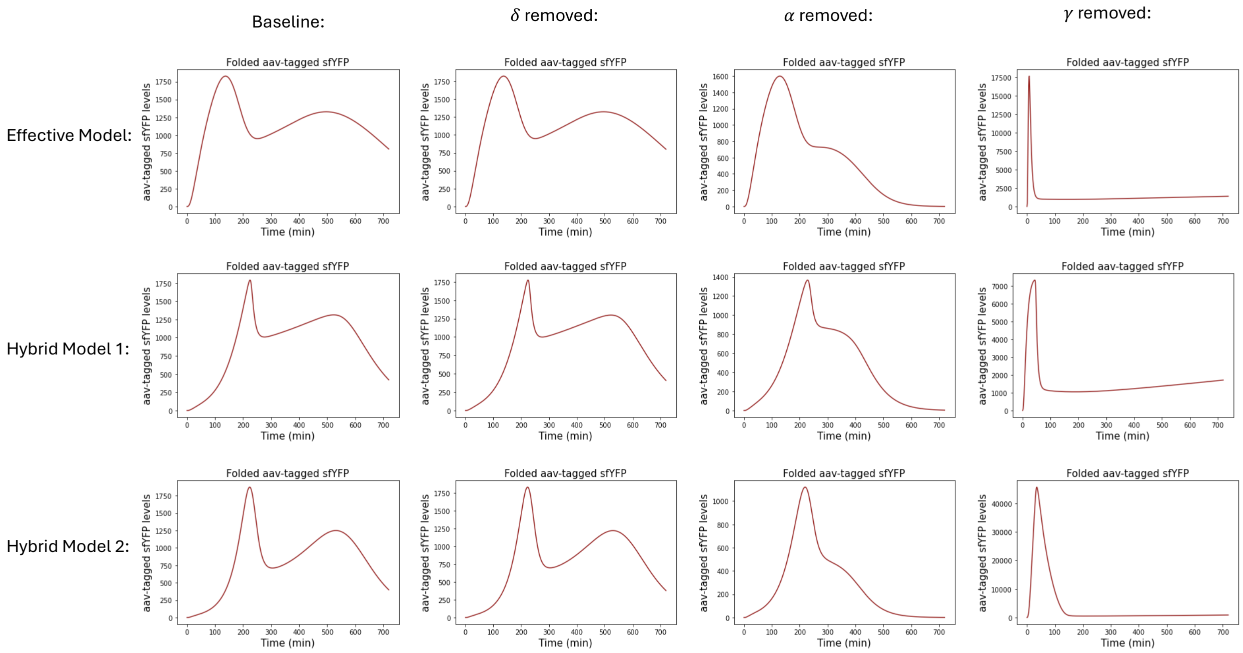


**Fig. S15.** Ablation study on the reduced models by systematically removing one RMF at a time

### S8 Layered Feedback Control Model

This section describes the effective model used to describe the dynamics of the layered feedback control construct (model used in text Figure 5). In this model, we have split the translation rate into a CRN, accounting for the resource allocation for translation. The parameters in bold indicate that they are modified by RMFs.

#### Ordinary Differential Equations

Inducer Cassette:

$$\frac{dM_{cin}}{dt} = \boldsymbol{\beta_A} \cdot \boldsymbol{\gamma_{tx}} \cdot \left(\frac{x}{K_x + x}\right) \cdot f_{trans} - (\boldsymbol{d_m} \cdot \boldsymbol{\alpha} + \boldsymbol{d_{dil}} \cdot \boldsymbol{\alpha}) \cdot M_{cin} - r_{intn_{cin}} + r_{elng_{cin}} \quad (S109)$$

$$\frac{dTI_{cin}}{dt} = r_{intn_{cin}} - r_{elng_{cin}} \quad (S110)$$

$$\frac{dP_{cin}}{dt} = r_{elng_{cin}} - (\boldsymbol{d} \cdot \boldsymbol{\delta} \cdot (\boldsymbol{1} - \boldsymbol{\psi_P}) + \boldsymbol{d_{dil}} \cdot \boldsymbol{\alpha}) \cdot P_{cin} - K_r \cdot P_{cin} \quad (S111)$$

$$\frac{dC_{cin}}{dt} = K_r \cdot P_{cin} - (\boldsymbol{d} \cdot \boldsymbol{\delta} \cdot (\boldsymbol{1} - \boldsymbol{\psi_P}) + \boldsymbol{d_{dil}} \cdot \boldsymbol{\alpha}) \cdot C_{cin} \quad (S112)$$

Reporter Cassette:

$$\frac{dM_G}{dt} = \boldsymbol{\beta_B} \cdot \boldsymbol{\gamma_{tx}} \cdot \left(\frac{C_{cin}}{K_{cin} + C_{cin}} + l_0\right) \cdot f_{cis} - (\boldsymbol{d_m} \cdot \boldsymbol{\alpha} + \boldsymbol{d_{dil}} \cdot \boldsymbol{\alpha}) \cdot M_G$$
$$-(r_{intn_{lac}} + r_{intn_{FP}}) + (r_{elng_{lac}} + r_{elng_{FP}}) \quad (S113)$$

$$\frac{dR}{dt} = \boldsymbol{\beta_B} \cdot \boldsymbol{\gamma} \cdot \left(\frac{C_{cin}}{K_{cin} + C_{cin}} + l_0\right) \cdot f_{cis} - (\boldsymbol{d_r} \cdot \boldsymbol{\alpha} + \boldsymbol{d_{dil}} \cdot \boldsymbol{\alpha}) \cdot R$$
$$-\boldsymbol{k_{fold_R}} \cdot (\boldsymbol{\gamma_{rate}} \cdot \boldsymbol{\psi_R} + \boldsymbol{b_{fold}}) \cdot R \quad (S114)$$

$$\frac{dR_m}{dt} = \boldsymbol{k_{fold_R}} \cdot (\boldsymbol{\gamma_{rate}} \cdot \boldsymbol{\psi_R} + \boldsymbol{b_{fold}}) \cdot R - (\boldsymbol{d_r} \cdot \boldsymbol{\alpha} + \boldsymbol{d_{dil}} \cdot \boldsymbol{\alpha}) \cdot R_m \quad (S115)$$

$$\frac{dTI_{lac}}{dt} = r_{intn_{lac}} - r_{elng_{FP}} \quad (S116)$$

$$\frac{dTI_{FP}}{dt} = r_{intn_{FP}} - r_{elng_{FP}} \quad (S117)$$

$$\frac{dP_{lac}}{dt} = r_{elng_{lac}} - (\boldsymbol{d} \cdot \boldsymbol{\delta} \cdot (\boldsymbol{1} - \boldsymbol{\psi_P}) + \boldsymbol{d_{dil}} \cdot \boldsymbol{\alpha}) \cdot P_{lac}$$
$$-\boldsymbol{k_{fold_{lac}}} \cdot (\boldsymbol{\gamma_{rate}} \cdot \boldsymbol{\psi_P} + \boldsymbol{b_{fold}}) \cdot P_{lac} \quad (S118)$$

$$\frac{dP_{m_{lac}}}{dt} = \boldsymbol{k_{fold_{lac}}} \cdot (\boldsymbol{\gamma_{rate}} \cdot \boldsymbol{\psi_P} + \boldsymbol{b_{fold}}) \cdot P_{lac} - (\boldsymbol{d} \cdot \boldsymbol{\delta} \cdot (\boldsymbol{1} - \boldsymbol{\psi_P}) + \boldsymbol{d_{dil}} \cdot \boldsymbol{\alpha}) \cdot P_{m_{lac}} \quad (S119)$$

$$\frac{dP_{FP}}{dt} = r_{elng_{FP}} - (\boldsymbol{d} \cdot \boldsymbol{\delta} \cdot (\boldsymbol{1} - \boldsymbol{\psi_P}) + \boldsymbol{d_{dil}} \cdot \boldsymbol{\alpha}) \cdot P_{FP}$$
$$-\boldsymbol{k_{fold_{FP}}} \cdot (\boldsymbol{\gamma_{rate}} \cdot \boldsymbol{\psi_P} + \boldsymbol{b_{fold}}) \cdot P_{FP} \quad (S120)$$

$$\frac{dP_{m_{FP}}}{dt} = \boldsymbol{k_{fold_{FP}}} \cdot (\boldsymbol{\gamma_{rate}} \cdot \boldsymbol{\psi_P} + \boldsymbol{b_{fold}}) \cdot P_{FP} - (\boldsymbol{d} \cdot \boldsymbol{\delta} \cdot (\boldsymbol{1} - \boldsymbol{\psi_P}) + \boldsymbol{d_{dil}} \cdot \boldsymbol{\alpha}) \cdot P_{m_{FP}} \quad (S121)$$

$$\frac{dC}{dt} = k_{gr} \cdot C \cdot \left(1 - \frac{C}{C_{max}}\right) \quad (S122)$$

Translation reactions:

$$r_{intn_{cin}} = k_{tli_{b_{ind}}} \cdot Rsrc_{free} \cdot M_{cin} - k_{tli_{u_{ind}}} \cdot TI_{cin} \quad (S123)$$
$$r_{elng_{cin}} = TI_{cin} \cdot k_{tl_{ind}} \quad (S124)$$
$$r_{intn_{lac}} = k_{tli_{b_{lac}}} \cdot Rsrc_{free} \cdot M_G - k_{tli_{u_{lac}}} \cdot TI_{lac} \quad (S125)$$
$$r_{elng_{lac}} = TI_{lac} \cdot k_{tl_{lac}} \quad (S126)$$
$$r_{intn_{FP}} = k_{tli_{b_{FP}}} \cdot Rsrc_{free} \cdot M_G - k_{tli_{u_{sFP}}} \cdot TI_{FP} \quad (S127)$$
$$r_{elng_{FP}} = TI_{FP} \cdot k_{tl_{FP}} \quad (S128)$$

Conservation equations for growth dependent resources tRNA ($T$) and Ribosome ($R$):

$$R_{rsrc_{total}} = R_{rsrc_{max}} \cdot \gamma_{rsrc}$$
$$R_{rsrc_{free}} = R_{rsrc_{total}} - TI_{cin} - TI_{lac} - TI_{FP} \quad (S129)$$

Feedback equations:

$$f_{cis} = \frac{K_R^{n_{cis}}}{K_R^{n_{cis}} + R_m^{n_{cis}}} \quad (S130)$$

$$f_{trans} = \frac{K_{lac}^{n_{trans}}}{K_{lac}^{n_{trans}} + P_{m_{lac}}^{n_{trans}}} \quad (S131)$$

**Table S10 GEAGS Layered Control Model Species**

| Species | Description |
|---|---|
| $x$ | Chemical inducer for first cassette |
| $M_{cin}$ | mRNA coding for CinR |
| $TI_{cin}$ | Translation initiation complex for CinR |
| $P_{cin}$ | Premature CinR |
| $C_{cin}$ | Mature CinR complex |
| $M_G$ | mRNA coding for LacI and sfYFP |
| $R$ | Unfolded sRNA |
| $R_m$ | Folded sRNA |
| $TI_{lac}$ | Translation initiation complex for LacI |
| $TI_{FP}$ | Translation initiation complex for sfYFP |
| $P_{lac}$ | Unfolded LacI |
| $P_{m_{lac}}$ | Mature LacI |
| $P_{FP}$ | Unfolded sfYFP |
| $P_{m_{FP}}$ | Mature sfYFP |
| $C$ | Cell count |

**Table S11 GEAGS Layered Control Model Parameters**

| Parameter | Description | Unit | Value |
|---|---|---|---|
| $\beta_A$ | Transcription rate per plasmid for inducer cassette | $nM \cdot min^{-1}$ | 0.14 |
| $K_x$ | Activation coefficient for chemical inducer | $nM$ | 1.4e4 |
| $K_R$ | Repression coefficient for sRNA | $nM$ | 45 |
| $K_{lac}$ | Repression constant for LacI | $nM$ | 153 |
| $d_m$ | mRNA degradation rate | $min^{-1}$ | 5.3e-2 |
| $k_{tl_{cin}}$ | Translation elongation rate of CinR | $min^{-1}$ | 3 |
| $d_p$ | Protein degradation rate | $min^{-1}$ | 9.9e-3 |
| $K_r$ | CinR maturation rate | $min^{-1}$ | 0.86 |
| $\beta_B$ | Transcription rate per plasmid for reporter cassette | $nM \cdot min^{-1}$ | 0.12 |
| $K_{cin}$ | Activation constant for CinR | $nM$ | 153 |
| $d_r$ | sRNA degradation rate | $min^{-1}$ | 5e-2 |
| $k_{fold_{FP}}$ | sfYFP folding rate | $min^{-1}$ | 0.14 |
| $k_{fold_{lac}}$ | LacI folding rate | $min^{-1}$ | 1e-2 |
| $k_{tl_{FP}}$ | Translation elongation rate of sfYFP | $min^{-1}$ | 0.2 |
| $k_{tl_{lac}}$ | Translation elongation rate of LacI | $min^{-1}$ | 2.78 |
| $k_{fold_R}$ | sRNA folding rate | $min^{-1}$ | 0.11 |
| $l_{0_{cin}}$ | Leak coefficient for $P_{cin}$ promoter | $N/A$ | 9.8e-2 |
| $R_{rsrc_{max}}$ | Maximum cap on translation resource | $nM$ | 420 |
| $k_{tlib_{lac}}$ | Translation initiation binding rate of LacI | $nM^{-1}min^{-1}$ | 9.9e-3 |
| $k_{tlib_{FP}}$ | Translation initiation binding rate of sfYFP | $nM^{-1}min^{-1}$ | 2.2e-2 |
| $k_{tlib_{cin}}$ | Translation initiation binding rate of CinR | $nM^{-1}min^{-1}$ | 7.1e-2 |
| $K_{tliu}$ | Translation initiation unbinding rate | $min^{-1}$ | 7 |
| $b_{fold}$ | Basal folding rate | $N/A$ | 0.1 |
| $K_{P_{total}}$ | Activation coefficient for RMF $\psi_P$ | $nM$ | 3.8e3 |

| | | | |
|---|---|---|---|
| $n_{\gamma_{rsrsc-tx}}$ | Exponent of $\gamma$ for transcription resources | $N/A$ | 0.25 |
| $n_{\gamma_{rsrsc-tl}}$ | Exponent of $\gamma$ for translation resources | $N/A$ | 0.4 |
| $n_{\gamma_{rate}}$ | Exponent of $\gamma$ for rates | $N/A$ | 6.5e-2 |
| $n_{Cin_{actn}}$ | Hill coefficient for $P_{Cin}$ activation | $N/A$ | 3 |
| $n_{Rhl_{actn}}$ | Hill coefficient for $P_{Rhl/Lac}$ activation | $N/A$ | 3 |
| $n_{\delta}$ | Hill coefficient for RMF $\delta$ | $N/A$ | 5.5 |
| $n_{\psi}$ | Hill coefficient for RMF $\psi_P$ | $N/A$ | 3 |
| $n_{trans}$ | Hill coefficient for trans repression | $N/A$ | 3 |
| $n_{cis}$ | Hill coefficient for cis repression | $N/A$ | 3 |
